# AI-Guided CAR Designs and AKT3 Degradation Synergize to Enhance Bispecific and Trispecific CAR-T Cell Persistence and Overcome Antigen Escape

**DOI:** 10.1101/2025.06.12.658477

**Authors:** Mohammad Sufyan Ansari, Varnit Chauhan, Aashi Singh, Areej Akhtar, Nisha Chaudhary, Reegina Tyagi, Divya, Kashif Husain, Sheetal Sharma, Ruquaiya Alam, Md Shakir, Mehak Pracha, Samreen Anjum, Mohd Nadeem, Md Imam Faizan, Iqbal Azmi, Aditya Ramdas Iyer, Pragya Gupta, Mehwish Nafiz, Shayan Ali, Insha Mohi Uddin, Momina Javid, Hamenth Kumar P, Amit Kumar Srivastava, Ulaganathan Mabalirajan, Vikram Mathews, Sivaprakash Ramalingam, Gaurav Kharya, Tanveer Ahmad

## Abstract

The structural design of chimeric antigen receptors (CARs) is critical for achieving robust and durable anti-tumor responses, particularly when targeting multiple antigens to prevent tumor antigen escape. However, increasing CAR complexity can introduce structural vulnerabilities, leading to antigen-independent T cell activation, activation-induced cell death, and reduced CAR-T cell persistence. To overcome these challenges, we designed 10,824 CAR molecules across diverse formats and screened 1,452 constructs in-vitro to develop an artificial intelligence model, termed CAR-Mediated Self-Destruction (CARMSeD), which predicts CAR designs susceptible to self-activation and dysfunction. Guided by CARMSeD and structural CAR-CAR interaction modeling, we identified optimized CAR architectures incorporating ICOS and 4-1BB co-stimulatory domains. Humanized bispecific CARs targeting CD20/CD19 and CD22/CD19 demonstrated superior anti-tumor efficacy and persistence both in-vitro and in various xenograft mice models. To further extend CAR-T cell persistence, we engineered bispecific CARs integrated with an AKT3-targeted PROTAC strategy. Targeted degradation of AKT3 enhanced anti-tumor potency, promoted memory T cell formation, and enabled sustained responses even under tumor rechallenge and CD19 antigen-loss conditions. Mechanistically, these effects were mediated by metabolic reprogramming involving FOXO4; notably, FOXO4-deficient CAR-T cells exhibited impaired long-term persistence. Leveraging these mechanistic insights, we developed a trispecific CAR-T cell platform incorporating a bispecific T cell engager (BiTE) targeting CD22/CD3, combined with AKT3 PROTACs. These trispecific CAR-T cells achieved potent tumor eradication, even against malignancies lacking both CD19 and CD20 expression. Collectively, this study presents a comprehensive strategy combining structure-based design, AI-guided screening, and targeted protein degradation to engineer next-generation bi and trispecific CAR-T cells with enhanced persistence, broad antigen coverage, and superior therapeutic durability.

## Introduction

CD19 CAR-T cell therapy has revolutionized the treatment of otherwise incurable B-cell malignancies; however, its long-term efficacy is frequently compromised by antigen escape, wherein malignant cells downregulate or lose CD19 expression to evade immune recognition^1–3^. This challenge is particularly pronounced in heterogeneous tumors containing antigen-low or antigen-negative subclones, often leading to relapse^2,4^. Our in-silico analyses, supported by multiple clinical datasets, consistently show that a significant proportion of patients relapse following CAR-T cell therapy due to modulation or complete loss of CD19 expression. To overcome this limitation, there has been increasing interest in dual and triple antigen targeting CAR-T cell strategies that simultaneously recognize CD19, CD20, and CD22^5–9^. These multi-antigen approaches aim to mitigate immune escape and offer broader tumor coverage. For instance, bispecific CAR-T cells targeting CD19 and CD20 have shown enhanced antitumor efficacy and reduced relapse in both preclinical models and clinical trials^10,11^. In a phase 1 study (NCT04007029), naïve/memory-derived CD19/CD20 CAR-T cells achieved a 90% overall response rate and 70% complete remission in patients with relapsed/refractory non-Hodgkin lymphoma, with low toxicity and durable outcomes^12^. Similarly, trispecific CAR-T cells targeting CD19, CD20, and CD22 have demonstrated the potential to overcome antigenic heterogeneity and improve long-term disease control^9,13^.

However, while these strategies address antigen escape, they introduce a new challenge of insufficient CAR-T cell persistence, particularly in complex multi-antigen designs. Although expanding antigen specificity reduces the risk of tumor escape, it often compromises T cell fitness and increases susceptibility to functional exhaustion^14,15^. This trade-off ultimately limits the sustained efficacy of treatment. As such, next-generation CAR-T engineering must develop integrated solutions that tackle both antigen escape and limited persistence in tandem CAR designs, which are comparatively better when translating into the clinical settings due to ease of manufacturing as compared to other approaches including co-transduction methods and bicistronic CAR designs^15,16^. Thus, in addition to immune evasion, limited CAR-T cell persistence itself remains a critical barrier to durable clinical outcomes. Although initial tumor clearance is often achieved, CAR-T cells frequently succumb to functional exhaustion, marked by impaired metabolic activity, reduced cytokine production, and poor memory T cell formation^17–20^. These factors are strongly associated with CD19-positive relapse over the years^21–23^. To combat this, co-stimulatory domains such as 4-1BB, ICOS and cytokines such as IL-12, IL-5 and IL-18 have been incorporated into CAR constructs to enhance T cell activation, longevity, and memory programming^24–28^. Third-generation CARs, which integrate multiple co-stimulatory domains, have also demonstrated superior persistence and antitumor function compared to earlier CAR designs^29,30^ Moreover, metabolic reprogramming strategies, particularly those enhancing mitochondrial oxidative phosphorylation (OXPHOS), have been shown to promote the formation of T stem cell memory (T_scm_), supporting long-term persistence and durable responses^31,32^.

A promising approach to boosting CAR-T cell persistence involves modulation of the AKT signaling pathway. AKT is a central regulator of cell metabolism, differentiation, and survival, and plays a critical role in T cell memory development^33–35^. Sustained AKT activation in CAR-T cells, particularly in exhausted states, impairs memory differentiation and favors terminal effector phenotypes. AKT and its downstream mTOR signaling regulate autophagy, mitophagy, and metabolic balance^36^. Inhibition of AKT or mTOR enhances memory-associated gene expression and supports metabolic fitness. Recent studies have shown that AKT suppresses FOXO1, a transcription factor essential for memory T cell development, by restricting its nuclear localization^37^. Inhibition of AKT restores FOXO1 activity, promoting T_scm_/T_cm_ differentiation, increased persistence, and enhanced antitumor function^35,38^. While the roles of AKT1 and AKT2 have been explored, the specific contribution of AKT3 in T cell memory remains uncharacterized. Similarly, among the four FOXO family members (FOXO1– 4), the functions of FOXO1–3 in memory T cell biology are better understood, while FOXO4 remains largely unknown in this context.

To address these knowledge gaps and improve CAR-T cell persistence, we developed a PROTAC-based approach to selectively degrade AKT3. Transcriptomic profiling identified AKT3 as differentially upregulated during CAR-T cell exhaustion. Targeted degradation of AKT3 using PROTACs resulted in enhanced T_em_/T_cm_ formation and metabolic fitness in multi-antigen targeting CAR-T cells. Notably, this effect was mediated through the upregulation of FOXO4, which emerged as a key regulator of memory differentiation and CAR-T persistence. This novel AKT3–FOXO4 axis provides a promising strategy for generating long-lasting, metabolically resilient CAR-T cells capable of overcoming both antigen escape and functional exhaustion.

## Results

### In silico analysis generated best-performing bispecific and trispecific CAR molecules

In an in-silico analysis of 4,129 patients who received CAR-T cell therapy, either monospecific or bispecific, we observed a significant relapse rate, consistent with findings from previous meta-analyses (**Figure 1A**, **Table 1 & S1A)** ^1–3^. Notably, a substantial proportion of patients treated with monospecific CAR-T cells experienced CD19-negative tumor recurrence (42.1%) (**Figure 1A**). Although prior studies have explored multi-antigen targeting CAR designs to mitigate antigen escape ^5–9^, these approaches still exhibited a high incidence of CD19 antigen loss (31.52%) and were associated with reduced persistence (**Figure S1B**). Importantly, the structural features of these multi-antigen CARs, and their potential contribution to activation induced cell death (AICD), have not been systematically investigated at the molecular level.

**Figure 1.**
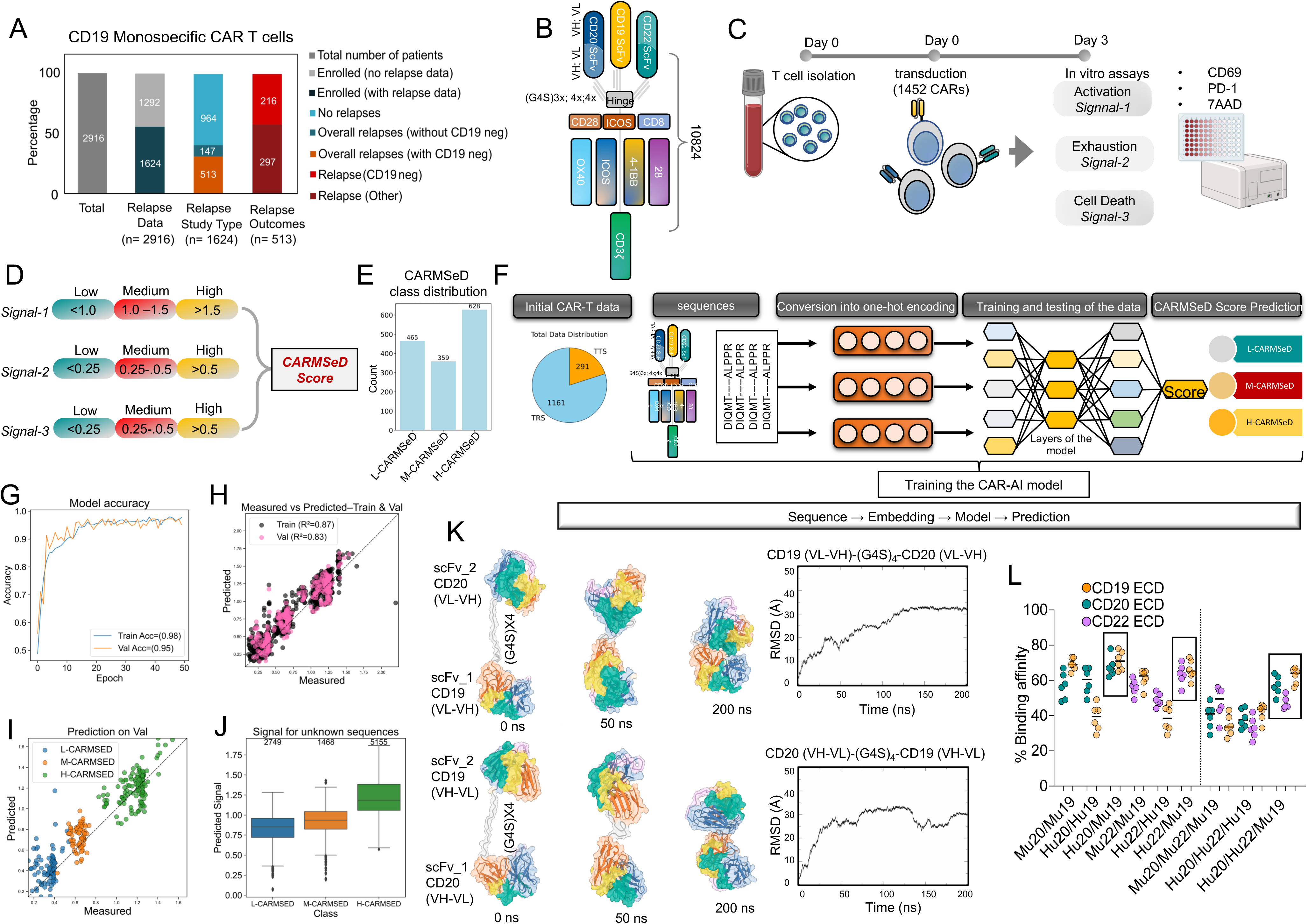
AI-guided screening and selection of optimized CAR constructs using CARMSeD scoring. (A) In-silico analysis of CAR-T cell-treated patients (n=4,219) revealed a high relapse rate, with 42.11% (n=216 of n=513 overall relapse patients) experiencing CD19-negative recurrence after monospecific CAR-Therapy (n=2,916). (B) Schematic overview of the CAR design strategy showing mono, bi, and trispecific constructs targeting CD19, CD20, and CD22. (C) Experimental workflow illustrating CAR screening: 1,452 CARs were transduced into primary T cells and analyzed for signal-1 (activation), signal-2 (exhaustion), and signal-3 (cell death). (D) Categorization of CARs into low (L), medium (M3), and high (H) levels based on fluorescence intensity cutoffs determined by monospecific CD19 CARs. (E) Distribution of 1,452 screened CARs across L-, M-, and H-CARMSeD categories using the CAR-Mediated Self-Destruction (CARMSeD) scoring system. (F) AI model development pipeline for CAR dysfunction risk prediction, based on 1,452 CAR constructs with an 80:20 split for training and testing. (G–J) Performance metrics of AI model predicting CAR-Mediated Self-Destruction (CAR-MSED) scores using 1,452 CAR constructs (G) Model accuracy over 50 epochs, achieving a training accuracy of 0.98 and validation accuracy of 0.95. (H) Scatter plot comparing measured versus predicted CAR-MSED scores for training (R^2^ = 0.87) and validation (R^2^ = 0.83) sets. (I) Predicted versus measured CAR-MSED scores on the validation set, categorized into low (L-CARMSED, blue), medium (M-CARMSED, orange), and high (H-CARMSED, green) groups. (J) Box plot of predicted signal scores for 9,372 unknown sequences, classified as L-CARMSED (2,749 sequences), M-CARMSED (1,468 sequences), and H-CARMSED (5,155 sequences). (K) Molecular dynamics simulation of CAR constructs with varying linker lengths, assessing CAR-CAR interaction. Structural conformations at 0 ns, 50 ns and 200 ns for different CAR scFv arrangements highlighting CDR regions (surface transparency 30%), Root Mean Square Deviation (RMSD) plots over 200 ns for the both constructs, respectively, indicating structural stability and conformational changes. (L) In vitro receptor binding affinity validation for top humanized scFvs of CD19, CD20, and CD22 CARs (n=6).

To overcome these structural constraints, we engineered CARs in mono-, bi-, and trispecific formats targeting CD19, CD20, and CD22 (**Figure 1B**). A comprehensive CAR library was created by varying G_4_S linker lengths, co-stimulatory domains (single or dual), and transmembrane domain configurations yielding a total of 10,824 distinct CAR constructs (**Table 2**). From this library, 1,452 CARs were selected for in-vitro screening (**Table 3**). These were introduced into T cells derived from three healthy donors and assessed for three parameters: T cell activation (signal-1), exhaustion (signal-2), and cell death (signal-3), using fluorescence-based assays with corresponding markers (**Figure 1C**).

Signal intensities were classified into low (L), medium (M), and high (H) categories (**Figure 1D**), with cutoff values established based on the response to CD19 stimulation by monospecific CARs (**Figure S2**). Based on a defined scoring system CARs were categorized into groups reflecting CAR-Mediated Self-Destruction (CARMSeD). Among the 1,452 screened CARs, 465 were classified as L-CARMSeD, 359 as M-CARMSeD, and 628 as H-CARMSeD (**Figure 1E**).

An AI model was developed using the data from these experiments (**Figure 1F**). The model was trained on 1,161 CARs, with the remaining used for validation, achieving a prediction accuracy of 95% (**Figure 1G-J**). We then applied this AI model to evaluate the entire library, identifying 2,749 CARs with low predicted CARMSeD scores. Among these, bispecific CD20/19 or CD22/19 CARs − incorporating ICOS transmembrane domains, a primary co-stimulatory domain, and 4-1BB as the secondary co-stimulatory domain, demonstrated optimal results. These top-performing constructs also shared a domain orientation of VH followed by VL (linked with 3 repeats G_4_S linkers), and scFvs separated by 4 repeat G_4_S linkers. These configurations were prioritized for further development.

As the scFvs were murine-derived, there was concern about immunogenicity due to human anti-mouse antibody (HAMA) responses, particularly when multiple murine scFvs are combined. To address this, we used an AI tool to humanize the scFvs and computationally evaluated receptor interactions (**Figure S3, S4 & Table 4**). We also developed a computational pipeline to determine the inter-CAR interaction that helped to find the CARs with least interaction for the bispecific and trispecific formats (**Figure 1K, Figure S5 and movies 1, 2**). Thus, based on the CARMSeD scoring and CAR-CAR interaction, we eventually chose the top performing humanized scFv of each CD19, CD20 and CD22 and further evaluated them for their binding affinity with their respective receptors using the in-vitro binding affinity analysis (**Figure 1L**). Ultimately, the two best-performing bispecific CARs and one trispecific CAR were selected for further validation.

### Preclinical validation of bispecific CAR-T cells targeting CD20/CD19 to overcome antigen escape

To evaluate the anti-tumor efficacy and in-vitro persistence of the selected CAR-T cell constructs, we first established a K562 cell line platform stably expressing CD19, CD20, or CD22 receptors − either individually, or all three simultaneously (**Figure 2A & Figure S6**). Co-culture of CAR-T cells with these engineered target cells demonstrated potent and specific anti-tumor responses across all antigen configurations (**Figure 2B**). Importantly, all tested CAR constructs exhibited robust anti-tumor activity in cells expressing either individual antigens or co-expressing multiple targets, surpassing the performance of a second-generation monospecific CD19 (m19) CAR containing a 4-1BB co-stimulatory domain (**Figure 2B**).

**Figure 2.**
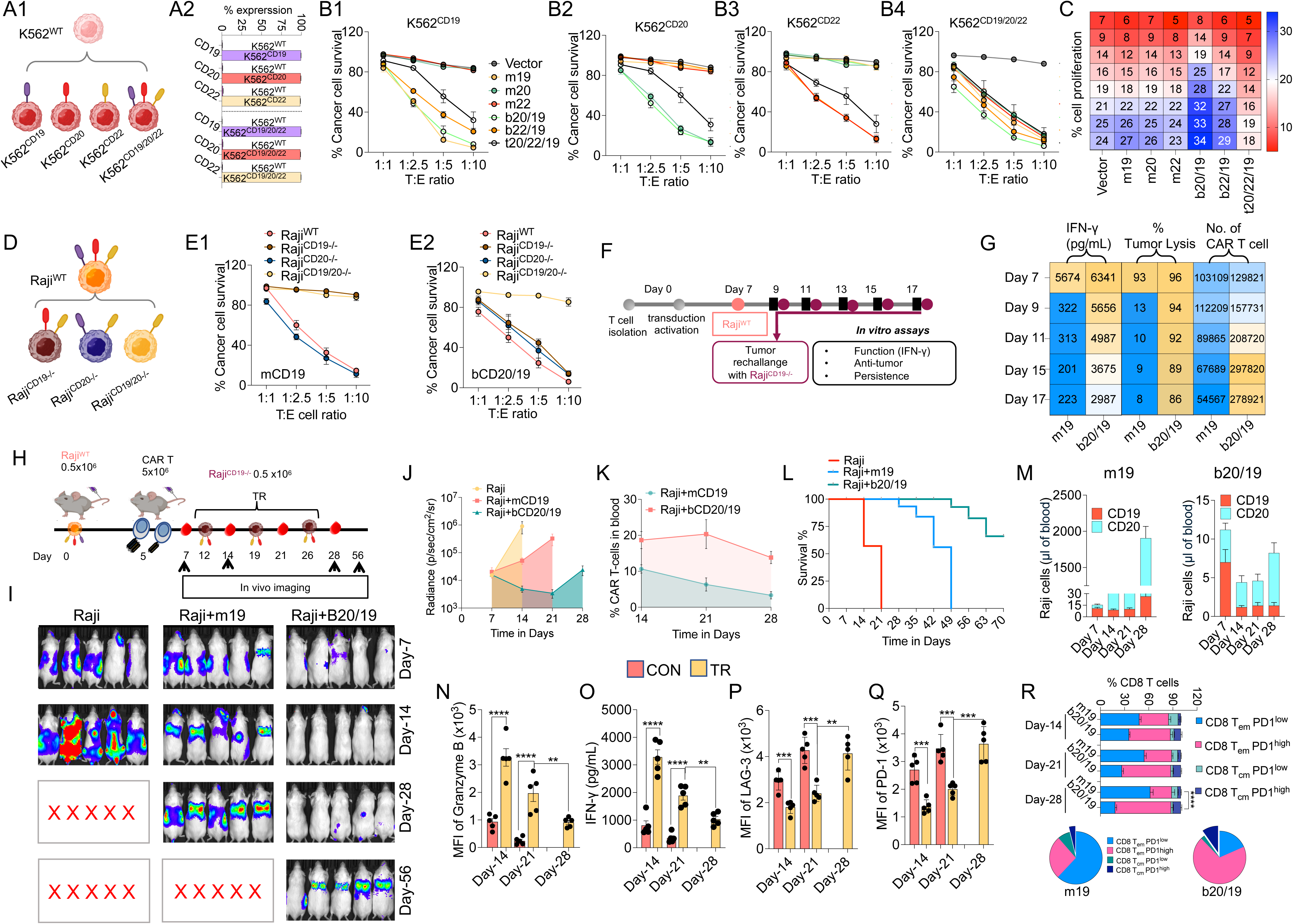
ICOS/4-1BB co-stimulatory bispecific CAR-T Cells prevent antigen escape in preclinical models. (A) Schematic illustration of the K562 cell line model expressing individual or triple combinations of CD19 (purple), CD20 (yellow), and CD22 (red) antigens. (B) Bar chart depicting the percentage expression of each antigen in K562 cell lines, both individually and in combination. (B) Cytotoxicity assays showing potent and antigen-specific killing of K562 target cells. All tested constructs surpassed the performance of second-generation monospecific CD19 (m19) CAR-T cells (n=5). (C) Comparison of proliferation rates for bispecific; b20/19 or b22/19, and trispecific; t20/19/22 CAR-T cells. Trispecific constructs showed reduced proliferation, consistent with increased structural rigidity predicted by CARMSeD scoring. (D) Schematic of the Raji WT cell line platform expressing CD19 (purple), CD20 (yellow), and CD22 (red) antigens, edited using CRISPR-Cas9 to generate knockout variants. (E) Cytotoxicity assays demonstrating the superior efficacy of b20/19 CAR-T cells in eliminating antigen-negative Raji variants, compared to ineffective m19 CARs (n=5). (F) Schematic representation of the tumor rechallenge (TR) model using the Raji WT cell line (Raji^WT^). Gray circles represent initial engraftment and monitoring phases, while purple circles indicate the timing of the Raji^CD19-/-^ rechallenge. (G) Heatmap representation of TR model showing IFN-γ secretion (pg/mL), percentage of tumor lysis, and the number of CAR-T cells detected on days 7, 9, 11, 15, and 17 post-rechallenge (n=5). (H) Schematic timeline of in vivo lymphoma model for evaluation of monospecific and bispecific CAR-T cells. Mice were xenografted with Raji^WT^ cells (expressing CD19, CD20, and CD22) (day 0), followed by administration of m19 or b20/19 CAR-T cells on day 5 and subsequent Raji^CD19-/-^ TR on day 12, 19 and 26. (I–J) Bioluminescent imaging and tumor burden quantification show effective tumor control by b20/19 CAR-T cells versus m19 CARs. (K) CAR-T cell persistence over time. (L) Kaplan-Meier survival curves showing survival outcomes over 70 days (n=5). (M) Analysis of residual tumor CD19 or CD20 tumor cells over time. (N, O) Bar plot showing Granzyme B and IFN-γ secretion from CD8^+^ CAR-T cells isolated post-treatment with m19 and b20/19 confirm functional cytotoxicity of b20/19 against CD19⁻ targets (n=5). (P–Q) Repeated TR induced upregulation of exhaustion markers PD-1 and LAG-3 (n=5). (R) Immunophenotyping of CAR-T cells post-TR shows loss of central memory (T_cm_) populations and increased PD-1 expression, consistent with functional exhaustion and limited persistence (n=5). Data represents mean ± SEM. *p < 0.05; **p < 0.01; ***p < 0.005; ****p < 0.001. A non-parametric t-test was used for statistical analysis between groups

To further delineate the functional impact of antigen targeting format on CAR-T cell expansion, we assessed the proliferative potential of bispecific (b20/19) and trispecific (t20/19/22) CAR-T cells. The trispecific construct exhibited reduced proliferation in comparison to the b20/19 configuration, aligning with computational predictions of increased structural rigidity and higher CARMSeD scores (**Figure 2C; Movies 3, 4**). Similarly, the b22/19 construct displayed a comparable proliferation rate to b20/19. Given its superior performance profile, we prioritized the b20/19 CAR-T cells for downstream in-vitro and in vivo studies.

To simulate clinical scenarios of antigen escape, we generated CD19^-/-^, CD20^-/-^ and CD19/20^-/-^ knockout variants of the Raji lymphoma cell line using CRISPR-Cas9 genome editing (**Figure 2D**). Cytotoxicity assays demonstrated that b20/19 CAR-T cells effectively eliminated both CD19^-/-^ and CD20^-/-^ Raji cells, whereas m19 CAR-T cells were ineffective against antigen-negative targets (**Figure 2E**). Building on previous findings^32^, we developed a tumor re-challenge (TR) model involving CD19^-/-^ Raji cells that retained CD20 expression upon re-introduction (**Figure 2F**). In this model, b20/19 CAR-T cells exhibited durable anti-tumor responses, sustained interferon-gamma (IFN-γ) production, and prolonged cellular persistence, as evidenced by bioluminescence imaging and functional assays (**Figure 2G**).

Following in-vitro validation, we transitioned to in vivo assessment using a xenograft mouse model rechallenged with CD19^-/-^ tumor cells (**Figure 2H**). Compared to the m19-treated group, b20/19 CAR-T cells achieved superior tumor clearance, reflected in a marked reduction of tumor bioluminescence signal and significant tumor regression (**Figure 2I, J**). However, tumor recurrence was observed over time. Longitudinal monitoring of peripheral blood samples from treated mice (n = 5 per group) indicated enhanced persistence of CAR-T cells in the b20/19 group (**Figure 2K**), which correlated with significantly improved survival outcomes (**Figure 2L**). Phenotypic characterization of residual tumor cells revealed a marked reduction in both CD19^+^ and CD20^+^ Raji cells in the b20/19 CAR-T cell-treated group. In contrast, mice treated with m19 CAR-T cells exhibited a progressive accumulation of CD20^+^ tumor cells, indicating selective antigen escape. Notably, the b20/19 group showed only a modest increase in CD20^+^ cells by Day 28, suggesting effective control of antigen escape mechanisms but lower persistence of these cells (**Figure 2M**).

Functional analysis of CD8^+^ CAR-T cells isolated from b20/19-treated mice revealed preserved effector function, as indicated by continued Granzyme B and IFN-γ secretion upon co-culture with CD19^-/-^ target cells (**Figure 2N, O**). However, with repeated tumor re-challenges, CAR-T cell function declined, concomitant with upregulation of exhaustion markers PD-1 and LAG-3. This phenotypic shift correlated with tumor relapse and a resurgence of CD20^+^ Raji cells (**Figure 2P, Q**). Immunophenotypic profiling further revealed a significant reduction in the central memory (T_cm_) CAR-T cell subset after repeated antigen exposure in the b20/19 group. This population showed elevated expression of PD-1, indicating progressive exhaustion and contributing to decreased persistence over time (**Figure 2R**).

### AKT3 is a key regulator of CAR-T cell metabolism, memory differentiation, and persistence

Limited persistence remains a critical barrier to the long-term success of current CAR-T cell therapies. Clinical data analysed shown in *Figure 1A* indicates that CAR-T cells fail to persist in 40–50% of patients, contributing to disease relapse and reduced therapeutic efficacy^1–3^. To address this limitation, we isolated CAR-T cells from control mice (CON; tumor only) and from an early tumor re-challenge (TR) group that had received three sequential tumor injections and b20/19 CAR-T cell therapy. Cells from each group (n=10) were pooled, and transcriptomic profiling via RNA sequencing was performed to identify differentially regulated genes and pathways (**Figure 3A**). The analysis revealed a significant dysregulation of genes associated with oxidative phosphorylation, mitochondrial metabolism, FOXO signaling and autophagy/mitophagy and genes implicated in T cell exhaustion (**Figure 3B**). Notably, AKT3 was among the most upregulated genes (**Figure 3C**), which was further confirmed at the protein level by flow cytometry (**Figure 3D**).

**Figure 3.**
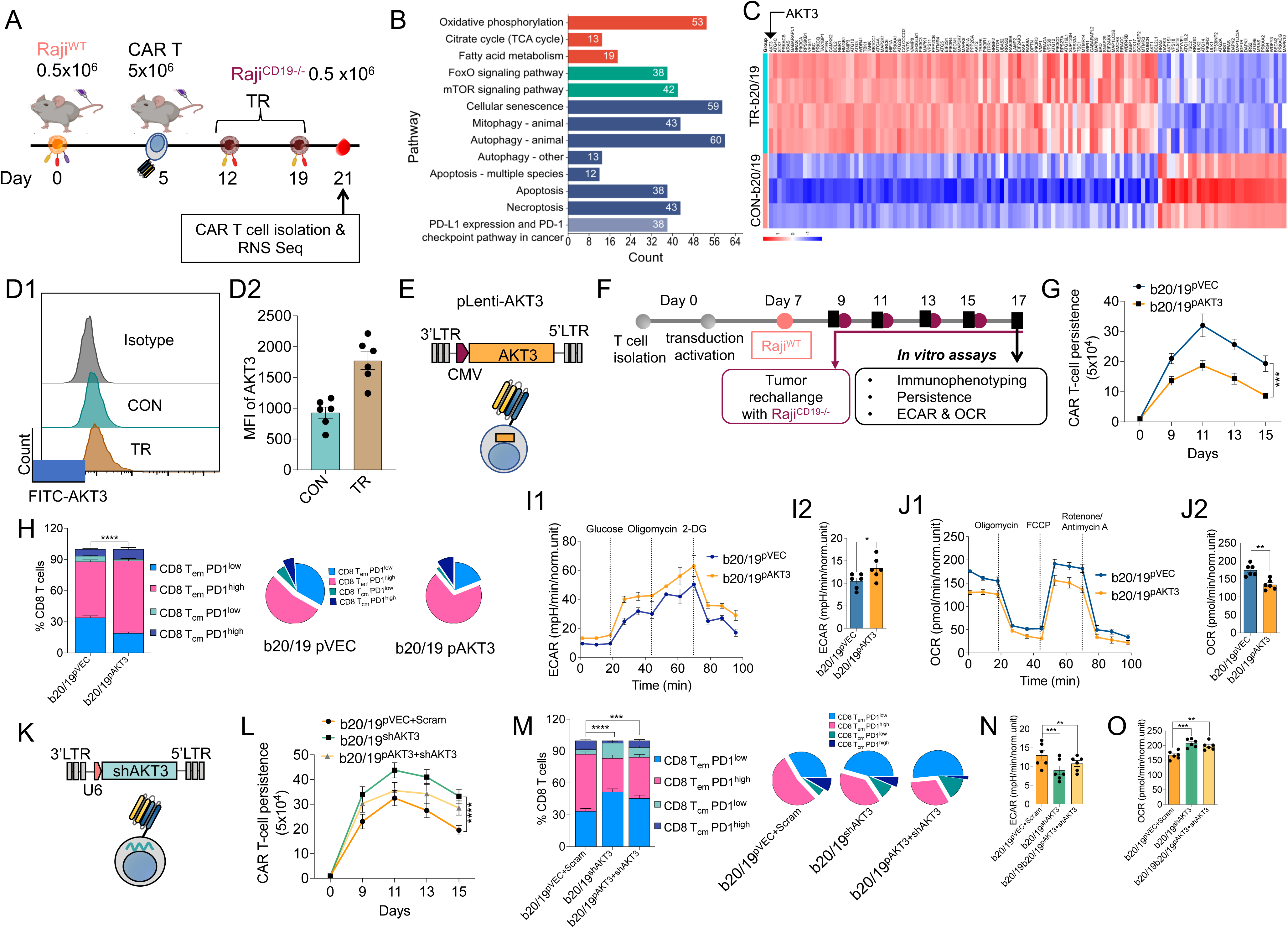
AKT3 upregulation in bispecific CAR-T cells limits persistence and promotes exhaustion. (A) Schematic timeline of in vivo lymphoma model for isolation of CAR-T cells for RNA seq analysis. (B) RNA seq showing pathway analysis of dysregulated gene associated with oxidative phosphorylation (OXPHOS), mitochondrial metabolism, FOXO signaling, and autophagy in TR group CAR-T cells. (C) Heatmap representing the comparative expression of dysregulated genes in autophagy highlighting AKT3 as one of the most significantly upregulated genes in TR group CAR-T cells. (D1) Representative flow cytometry histogram showing the fluorescence intensity of FITC-labeled AKT3 (FITC-AKT3) in T cells from different groups. (D2) Bar graph showing the mean fluorescence intensity (MFI) of AKT3 in T cells from the CON and TR groups (n=6). (E) Illustration of the plasmids showing pLenti-AKT3 lentiviral vector used for co-transduction of CAR-T cells with CAR vectors. (F-H) Schematic representation of the 17-day in vitro experimental timeline for evaluating T cell responses in TR model. T cells are isolated and activated from Day 0 to Day 7, followed by tumor challenge with Raji^WT^ cells on Day 7. (G) Line graph comparing the persistence of CAR-T cells transduced with b20/19 CAR and either a control vector (b20/19^pVEC^, blue circles) or an AKT3-expressing vector (b20/19^pAKT3^, orange squares) over time. (H) Histogram analysis and pie chart showing distribution of T_em_ and T_cm_ with PD-1^low^ or PD-1^high^ within these subsets (n=5). (I1) Line graph showing Extracellular Acidification Rate (ECAR, mpH/min) over 100 minutes. (I2) Bar graph of basal ECAR (mpH/min, normalized) between b20/19^pVEC^ (blue) and b20/19^pAKT3^ (orange) groups (n=6). (J1, J2) Similarly line graph showing Oxygen Consumption Rate (OCR, pMoles/min) and corresponding bar graph (n=6). (K) Schematic of the lentiviral vector delivering shRNA against AKT3 (shAKT3). The lower panel depicts the lentiviral particle transducing target CAR-T cells. (L) Line graph comparing the persistence of CAR-T cells over 15 days for three groups: b20/19^pVEC^ ^+ Scram^ (orange circles, control with scrambled shRNA), b20/19^shAKT3^ (green squares, AKT3 knockdown), and b20/19^pAKT3^ ^+ shAKT3^ (yellow triangles, AKT3 overexpression with knockdown) (n=5). (M) Bar graph and pie charts showing the percentage of various subsets of CD8^+^ CAR-T cells across three groups (n=5). AKT3 knockdown (b20/19^shAKT3^) significantly increases PD-1^low^ subsets, indicating decreased exhaustion after knowdown of AKT3. (N–O) Bar graph of ECAR, and OCR across three groups (n=6). Data represents mean ± SEM. *p < 0.05; **p < 0.01; ***p < 0.005; ****p < 0.001. A non-parametric t-test was used for statistical analysis between groups

To explore the functional implications of AKT3 overexpression, we engineered CAR-T cells co-transduced with a lentiviral vector encoding AKT3 (**Figure 3E & S7A**) and assessed their performance in the in-vitro tumor rechallenge (TR) model (**Figure 3F**). AKT3-overexpressing CAR-T cells exhibited a marked reduction in persistence over time (**Figure 3G**). Immunophenotyping revealed a decrease in T_cm_ and T_em_ subsets within PD-1^low^ populations, with a concomitant expansion of PD-1^high^ exhausted-like subsets (**Figure 3H**). Given the known role of AKT3 in cellular metabolism, we examined metabolic activity using ECAR and OCR assays. AKT3 overexpression led to a significant decrease in OXPHOS and a compensatory increase in glycolytic activity (**Figure 3I, J**).

To further investigate the causal role of AKT3, we employed shRNA-mediated knockdown of AKT3 in CAR-T cells (**Figure 3K & S7B**). In contrast to the overexpression model, AKT3 knockdown significantly enhanced long-term CAR-T cell persistence and increased the proportion of T_cm_ subsets within the PD-1^low^ population (**Figure 3M**). Metabolic profiling of these cells showed restored OXPHOS activity and reduced glycolysis, indicating a metabolic reprogramming favorable for memory cell formation (**Figure 3N, O**). Together, these findings establish AKT3 as a key regulator of CAR-T cell metabolic state, memory differentiation, and persistence. Targeting AKT3 may offer a promising strategy to enhance the durability and efficacy of CAR-T cell therapies.

### AKT3 PROTAC bispecific CAR-T cells demonstrated higher memory CAR-T cells

AKT3 inhibition has previously been associated with the induction of a memory phenotype in T cells. Recent studies underscore its role in enhancing effector function and long-lasting memory formation in CAR-T cells^35,38^. However, existing strategies rely heavily on ex vivo application of molecular inhibitors during CAR-T cell culture. To circumvent the need for such external supplementation, we developed a novel design to achieve intracellular AKT3 inhibition using a PROTAC-based approach. We first generated a series of AKT3-targeting peptides using RF diffusion modeling and validated them in silico (**Figure 4A & S8**). The top five candidates were selected for in-vitro screening. These peptides were fused to a cell-penetrating peptide (CPP) and tested in HEK-293T cells expressing a GFP-AKT3-mCherry reporter (**Figure S9A**). All peptides demonstrated dose-dependent AKT3 degradation, with peptide 2 (P2) exhibiting the strongest effect, confirmed by immunoblotting across multiple concentrations (**Figure S9B, C**). For functional validation in CAR-T cells, b20/19 CAR-T co-transduced with GFP were incubated with various concentrations of P2, resulting in a consistent, dose-dependent reduction of the endogenous AKT3 levels (**Figure 4B, C**). A non-targeting peptide (NTP) control showed no significant effect (**Figure 4C, D**). Subsequently, we engineered b20/19 CAR constructs incorporating the AKT3-PROTAC module and evaluated their function in CAR-T cells challenged with CD19^-/-^ tumor cells (**Figure 4E**). Upon repeated tumor rechallenge, fluorescence-based analysis confirmed AKT3 degradation specifically in PROTAC-modified CAR-T cells, while NTP-treated cells retained AKT3 expression (**Figure 4F**). These findings were validated by immunoblot and immunofluorescence, confirming endogenous AKT3 proteolysis (**Figure 4G, H**).

**Figure 4.**
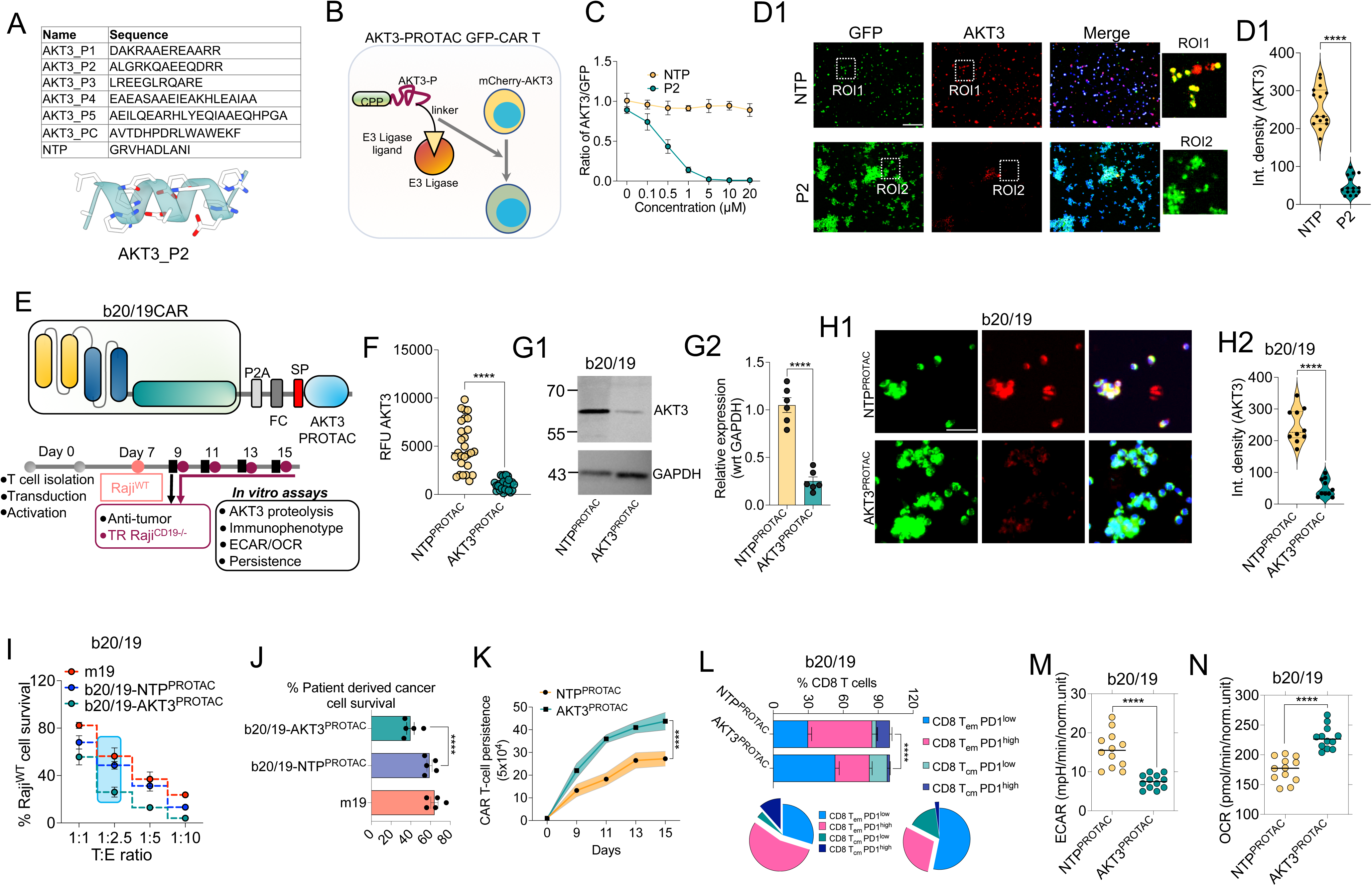
AKT3 PROTAC-engineered bispecific CAR-T cells exhibit enhanced memory phenotype and persistence. (A) Table listing the sequences of peptides designed using R.F. diffusion for PROTAC application, targeting AKT3 (AKT3_P1 to AKT3_P5, AKT3_PC) and a non-targeting peptide (NTP), with AKT3_P2 highlighted as the selected peptide. (B) Schematic illustrating the mCherry-AKT3 CAR-T cells induced with PROTAC. The PROTAC consists of the AKT3-targeting peptide (AKT3-P) fused to a cell-penetrating peptide (CPP) and via a linker connected to an E3 ligase ligand, enabling targeted degradation of AKT3 in GFP expressing CAR-T cells through the E3 ligase pathway. The AKT3 was detected using anti-AKT3 antibody (C) Line graph showing the dose-dependent effect of PROTAC peptide P2 (blue circles) compared to a non-targeting peptide (NTP, orange squares) on the ratio of mCherry to GFP fluorescence in CAR-T cells (n-4). (D1) Representative fluorescence microscopy images of CAR-T cells treated with NTP or P2, showing GFP (green), mCherry-AKT3 (red), and merged channels with regions of interest (ROIs). ROIs highlight reduced mCherry-AKT3 signal in P2-treated cells compared to NTP-treated cells. (D2) Histogram comparing the integrated density of AKT3 in NTP-treated (blue) and P2-treated (black) CAR-T cells (n=15 images). (E) Schematic of the b20/19 CAR construct, including a signal peptide (SP), furin cleavage (FC), and AKT3 PROTAC (P2A linker), designed to target AKT3 for degradation in CAR-T cells. Timeline of the 15-day experiment for evaluating b20/19 CAR-T cell responses. (F) Graph showing the relative fluorescence units (RFU) of AKT3 in b20/19 CAR-T cells treated with NTP or AKT3 PROTAC (n=25 data points). (G1) Western blot analysis of b20/19 CAR-T cells treated with NTP^PROTAC^ or AKT3^PROTAC^, probing for AKT3, and GAPDH as a loading control (G2) Bar graph showing the densitometry analysis of the blots (n=6). (H1) Representative fluorescence microscopy images of b20/19 CAR-T cells treated with non-targeting peptide (NTP^PROTAC^) or AKT3^PROTAC^, showing GFP (green), mCherry-AKT3 (red), and merged channels. (H2) Histogram comparing the integrated density of AKT3 in NTP^PROTAC^ and AKT3^PROTAC^ treated b20/19 CAR-T cells (n=10). (I) Percentage of Raji^WT^ cell survival after co-culture with CAR-T cells at different target-to-effector (T:E) ratios (n=4). (J) Percentage of patient-derived cancer cell survival after 24-hour co-culture with CAR-T cells at a target-to-effector (T:E) ratio of 1:2.5. b20/19-AKT3^PROTAC^ exhibits significantly higher cytotoxicity in patient derived cells (n=5). (K) In vitro tumor rechallenge model showing persistence of b20/19-AKT3^PROTAC^ CAR-T cells over time (n=4). (L) Flow cytometry histogram showing various T cell subsets (n=5). (M–N) Metabolic profiling showing ECAR and OCR analysis (n=12 data points). Data represents mean ± SEM. ****p < 0.001. A non-parametric t-test was used for statistical analysis between groups

By day 7 post-transduction, b20/19 AKT3^PROTAC^ CAR-T cells demonstrated potent cytotoxic activity across multiple CD19^+^ lymphoma and leukemia cell lines, as well as in patient-derived leukemia samples (**Figure S10A, B**). Remarkably, even at half the dose, b20/19-AKT3^PROTAC^ CARs maintained equivalent anti-tumor efficacy compared to mCD19 and b20/19 CARs (**Figure 4I**) and were also effective against CD19^-/-^ Raji cells (**Figure S10C**). Strong anti-tumor activity was observed in patient-derived leukemia models as well (**Figure 4J**). In-vitro TR modeling showed significantly enhanced persistence of b20/19-AKT3^PROTAC^ CAR-T cells (**Figure 4K**). Immunophenotyping revealed increased frequencies of T_cm_ and T_em_ subsets with reduced exhaustion markers (**Figure 4L**). Metabolic analysis further supported these observations, showing elevated OXPHOS and reduced reliance on glycolysis (**Figure 4M, N**). These results establish a robust, targeted approach to degrade AKT3 in CAR-T cells via PROTAC technology, enhancing memory phenotype, persistence, and anti-tumor efficacy, which was further validated in patient-derived tumor samples.

### AKT3 PROTACs alleviate CAR-T dysfunction by increasing FOXO4 expression and inhibiting mTOR activity

To elucidate the molecular basis by which AKT3 modulation enhances memory T cell formation, we investigated potential downstream targets of AKT3 (**Figure 5A**). CAR-T cells undergoing TR were harvested for mRNA extraction, and the expression of proteins associated with AKT3 was analyzed at the transcript level. Notably, b20/19 AKT3^PROTAC^ CAR-T cells displayed a significant upregulation of FOXO4 mRNA following TR (**Figure 5B**). This transcriptional increase was further confirmed at the total protein and phosphorylated protein levels (**Figure 5C**). To determine whether FOXO4 mediates the observed memory phenotype, we performed shRNA-mediated knockdown of FOXO4 (**Figure S11**). FOXO4 knockdown resulted in a marked increase in PD-1^high^ T memory subsets and a concomitant decrease in PD-1^low^ memory cells (**Figure 5D**). Functionally, this shift was associated with reduced CAR-T cell persistence (**Figure 5E**).

**Figure 5.**
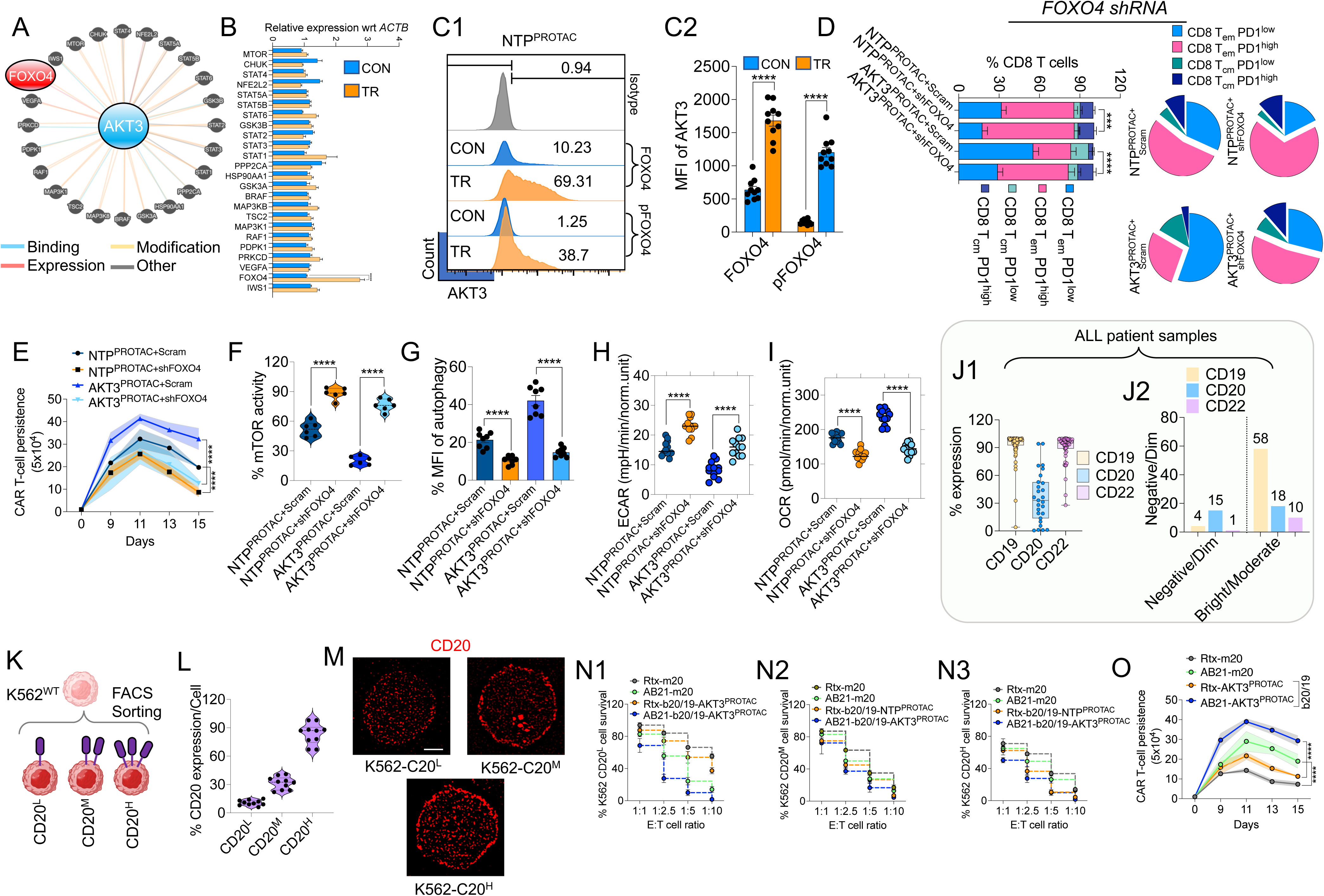
AKT3 PROTACs enhance CAR-T function by upregulating FOXO4 and suppressing mTOR signaling. (A) Pathway analysis of proteins involved in AKT3 interaction, modifications or regulation of its expression with emphasis on FOXO4. (B) Relative mRNA expression levels (normalized to beta actin) of key genes show upregulation of FOXO4 mRNA in b20/19-AKT3^PROTAC^ CAR-T. (C1) Flow cytometry histograms of total FOXO4 and phosphorylated FOXO4 (p-FOXO4) in CAR-T cells after TR with Raji^CD19-/-^ cells (C2) Histogram analysis of the flow cytometry plots (n=10). (D) Bar graph shows the percentage of CD8^+^ CAR-T cells expressing different phenotypes. Pie charts illustrate the proportional distribution of these subsets across conditions. (E) Persistence of CAR-T cells over 15 days under various conditions (n=4). (F) Violin plots show the percentage of mTOR activity (% mTOR activity) in various conditions, with shRNA based FOXO4 knockdown significantly elevated mTOR activity (n=6). (G) Bar plots show the percentage of MFI of autophagy from autophagic flux assay (n=8). (H) ECAR in NTP^PROTAC+Scram^, NTP^PROTAC+shFOXO4^, AKT3^PROTAC+Scram^, and AKT3^PROTAC+shFOXO4^ conditions, with FOXO4 knockdown increasing shift from oxidative phosphorylation (OXPHOS) to glycolysis (n=12 data points). (I) Similarly, OCR with FOXO4 knockdown decreasing mitochondrial respiration. Individual data points are shown for each condition (n=12 data points). (J1) Percentage of expression (% expression) of CD19 (yellow), CD20 (blue), and CD22 (purple) across 129 ALL patient samples, with varying expression levels for each marker. (J2) Bar graph displays the number of patient samples categorized as Negative/Dim, Moderate, or Bright for CD19, CD20, and CD22 expression. (K) Schematic illustration of K562^WT^ cells based on CD20 expression levels, resulting in three populations: CD20^L^ (low), CD20^M^ (medium), and CD20^H^ (high). (L) Violin plots show the percentage of CD20 expression (% CD20 expression) in the sorted K562^WT^ cell populations, confirming distinct expression levels (n=10). (M) Representative super-resolution microscopy images of differential CD20 surface expression in K562 cells. Images show DAPI (blue, nuclear staining) and CD20 (red) in K562-C20^L^ (low), K562-C20^M^ (medium), and K562-C20^H^ (high) cell. Scale bar indicates 10 μm. (N) Survival of K562 cells expressing varying CD20 expression levels under CAR-T cell treatments. Panels N1 (K562-CD20^L^), N2 (K562-CD20^M^), and N3 (K562-CD20^H^) show the percentage of CD20^+^ cell survival when treated with Rituximab-based monospecific CAR (Rtx-m20, dark green), in-house humanized anti-CD20 CAR (AB21-m20, green) (N=4). (O) Persistence of CAR-T cells with varying CD20-targeting CAR constructs over 15 days (N=5). Data represents mean ± SEM. ****p < 0.001. A non-parametric t-test was used for statistical analysis between groups. Scale bar indicates 10 μm.

Given the role of FOXO4 in autophagy via suppression of mTOR signaling, we next assessed mTOR activity. FOXO4 knockdown led to increased mTOR activation and significantly impaired autophagy (**Figure 5F, G**). Metabolic profiling further revealed a shift from OXPHOS to glycolysis in FOXO4-deficient cells (**Figure 5H, I**). These findings suggest that AKT3 tightly regulates FOXO4, possibly through phosphorylation and transcriptional repression, although further mechanistic studies are warranted. Overall, our data indicate that AKT3 proteolysis via PROTACs enhances CAR-T cell function and long-term persistence by promoting FOXO4-mediated autophagy and metabolic fitness.

Unlike CD19, which is robustly and uniformly expressed, CD20 displays heterogeneous and often dim surface expression. Analysis of clinical data from over 129 patients confirmed that CD20 expression is variable and frequently low (**Figure 5J & Table 5**). To model this heterogeneity, we engineered K562 cells expressing low, medium, and high levels of CD20 (**Figure 5K**) and validated single-cell expression by flow cytometry (**Figure 5L**). Immunofluorescence and super-resolution microscopy further confirmed differences in CD20 surface density (**Figure 5M & S12**). We next evaluated the anti-tumor activity of b20/19 AKT3^PROTAC^ CAR-T cells against these engineered targets. Compared to standard CD20-specific CAR constructs used in clinical trials, our optimized humanized CD20 scFv combined with AKT3-PROTAC demonstrated robust cytotoxicity even against cells with very low CD20 expression (**Figure 5N**). Moreover, b20/19-AKT3^PROTAC^ CAR-T cells exhibited superior persistence when challenged with CD20^low^ targets (**Figure 5O**), highlighting their potential for prolonged and effective tumor control even in settings of antigen heterogeneity.

### Higher In Vivo Persistence of AKT3-Targeted PROTAC CAR-T Cells

To determine the in vivo efficacy of b20/19-AKT3^PROTAC^ CAR-T cells, we developed a xenograft mouse model (**Figure 6A**). Mice were monitored longitudinally for tumor clearance and survival outcomes. Notably, mice treated with b20/19-AKT3^PROTAC^ CAR-T cells exhibited significantly superior tumor control and enhanced survival compared to controls (**Figure 6B–D**). In contrast, animals treated with CAR-T cells incorporating shRNA-mediated FOXO4 knockdown did not show comparable anti-tumor activity, emphasizing the critical role of FOXO4 in mediating therapeutic benefit. Importantly, longitudinal tracking revealed durable CAR-T cell persistence in the b20/19-AKT3^PROTA^C group, with detectable CAR-T cells up to 84 days post-infusion—the latest timepoint assessed (**Figure 6E**). Consistent with tumor clearance, no residual Raji cells were detected in these animals when evaluated at day 56 (**Figure 6F**). Further phenotypic analysis of CAR-T cells isolated 28 days post-infusion showed an increased proportion of memory T cell subsets and reduced expression of exhaustion markers (**Figure 6G**). Metabolic profiling revealed a predominant reliance on OXPHOS over glycolysis in b20/19-AKT3^PROTAC^ CAR-T cells, supporting a metabolically favorable memory phenotype (**Figure 6H, I**).

**Figure 6.**
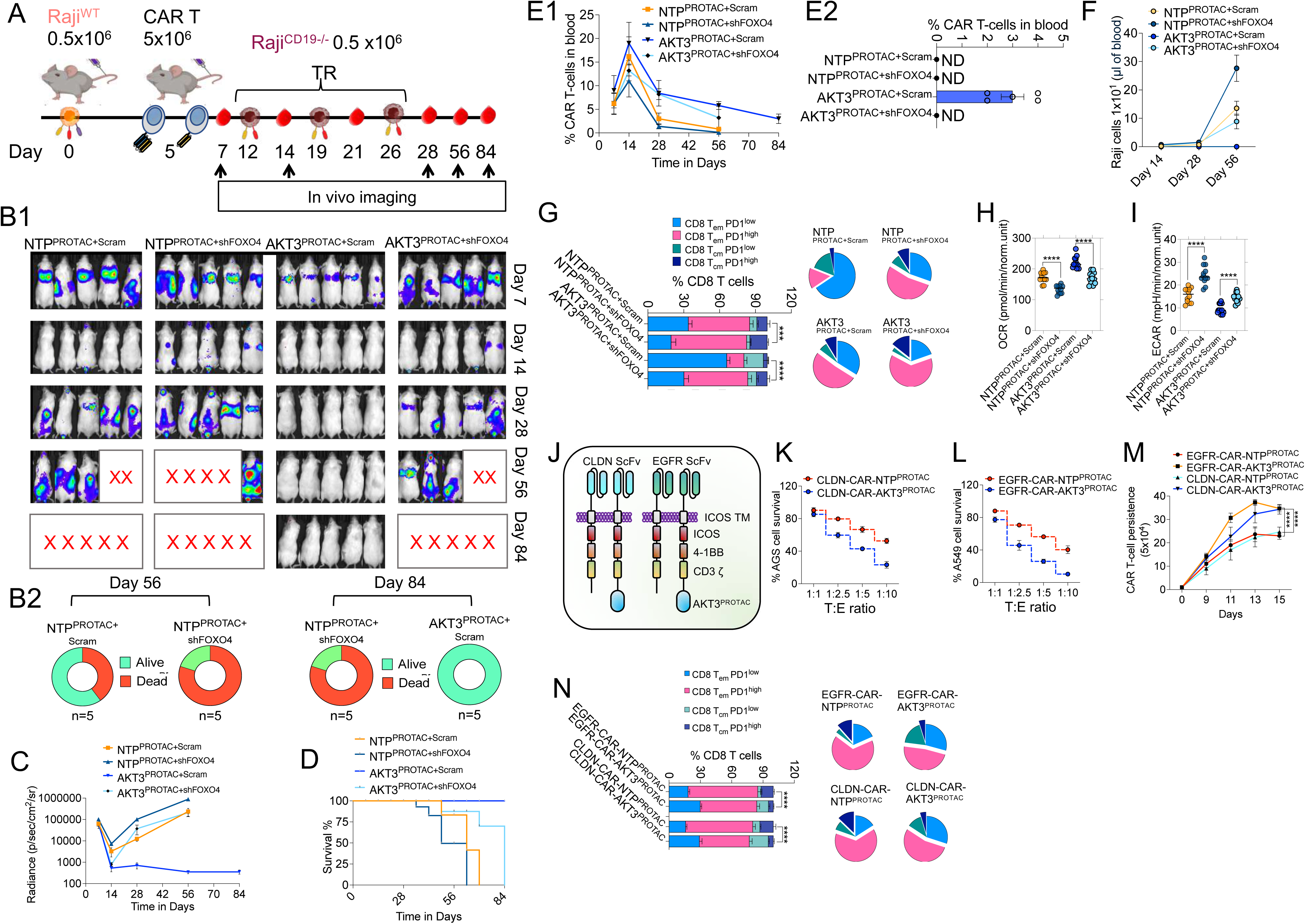
Higher in vivo persistence and broad efficacy of AKT3-targeted PROTAC CAR-T cells. (A) Schematic timeline of the experiment showing Raji^WT^ cell injection, CAR-T cell administration, and Raji^CD19-/-^ TR. (B1) In vivo bioluminescence imaging of mice treated with NTP^PROTAC+Scram^, NTP^PROTAC+shFOXO4^, AKT3^PROTAC+Scram^, and AKT3^PROTAC+shFOXO4^, showing tumor burden (red indicates high tumor load, blue indicates low) over 84 days, with ‘X’ marking deceased mice. (B2) Pie charts depict survival outcomes on days 56 and 84 (n=5). (C) Tumor radiance over 84 days, demonstrating reduced tumor burden in the AKT3^PROTAC+scram^ group. (D) Kaplan-Meier survival curves, with the AKT3^PROTAC+scram^ group exhibiting the highest survival rate. (E1) Line graph shows the percentage of CAR-T cells in blood (% CAR-T cells in blood) over 84 days. (E2) Bar graph displays the percentage of CAR-T cells in blood at day 84, with the AKT3^PROTAC+scram^ group showing detectable levels (∼3%), while other groups show non-detectable (ND) levels (n=5). (F) Tumor burden assessment till day 56. The line graph shows the number of Raji cells over time. The AKT3^PROTAC+scram^ group exhibits no detectable Raji cell burden by day 56, while other groups show some detectable cells (n=5). (G) Bar graph shows the percentage of CD8 CAR-T cells (% CD8 T cells) expressing different phenotypes on day 28 post-infusion with corresponding pie charts illustrating the proportional distribution (n=5). (H-I) ECAR and OCR under various conditions on day 28 (n=12 data points). (J) Schematic of CLDN ScFv and EGFR ScFv CAR constructs with ICOS, 4-1BB, CD3ζ, and AKT3^PROTAC^ domains. (K-L) Cytotoxicity of CLDN-CAR and EGFR-CAR-T cells with NTP3^PROTAC^ or AKT3^PROTAC^ against AGS (K) and A549 (L) cells at varying T:E ratios (n=5). (M) CAR-T cell proliferation over 15 days, showing enhanced expansion with AKT3^PROTAC^. (N) Bar graph and pie charts analysis of EGFR-CAR and CLDN-CAR-T cells with NTP^PROTAC^ or AKT3^PROTAC^, showing the percentage of various T cell subsets (n=5). Data represents mean ± SEM. ***p < 0.005; ****p < 0.001. A non-parametric t-test was used for statistical analysis between groups.

To test the robustness of the therapy, we reduced the administered CAR-T cell dose by half. Remarkably, similar tumor clearance and survival kinetics were observed even at the lower dose (**Figure S13**), underscoring the potency and persistence of AKT3-targeted PROTAC CAR-T cells.

To explore whether this strategy could be extended to solid tumors, we next engineered PROTAC-CAR-T cells targeting Claudin18.2 (CLDN; for gastric cancer) and EGFR (for non-small cell lung carcinoma; NSCLC) (**Figure 6J**). Compared to conventional CAR designs, PROTAC-engineered CAR-T cells exhibited significantly enhanced anti-tumor activity against gastric and NSCLC tumor models, respectively (**Figure 6K, L**). Notably, higher CAR-T cell persistence was again observed in the PROTAC groups, corroborated by an increased frequency of memory T cell subsets and reduced exhaustion status (**Figure 6M, N**). These findings highlight the broad applicability of PROTAC-based AKT3 targeting as a powerful strategy to enhance CAR-T cell persistence, functionality, and durability, not only in hematologic malignancies but also in solid tumor settings.

### Trispecific CAR exhibits safety and sustained cellular persistence

To further increase the versatility of the antibodies with three antigens, we made a strategy of using the b20/19AKT3^PROTAC^ with CD22 secretory BITE based on the CD3 and CD22 nanobody (nb) **(Figure 7A)**. The humanised nanobodies were designed using the computational approach and the CDR grafting method as described in *methods*. Initially to demonstrate the expression of nbCD3/22, we transduced the LVV encoding the b20/19AKT3^PROTAC^ plasmid encoding nbCD3/22 (hereafter b20/19AKT3^PROTAC+nbCD3/22^) in T cells and evaluated the expression on day 7 after transduction (**Figure S14A**). A dose dependent increase in the detection of nbCD3 and nbCD22 was observed in the culture supernatant detected using the fluorescent ELISA assay for nbCD3/22 (**Figure S14B, C**). To assess the relationship between BITE expression and CAR expression, we inhibited BITE secretion using Brefeldin A and performed intracellular flow cytometry analysis. CD22-PE and CD3-FITC antibodies were used to detect intracellular nbCD22 and nbCD3, respectively. Results demonstrated that nanobody expression positively correlated with surface expression of CD19 and CD20 CARs on T cells (**Figure 7B**), indicating coordinated expression and secretion of the engineered constructs.

**Figure 7.**
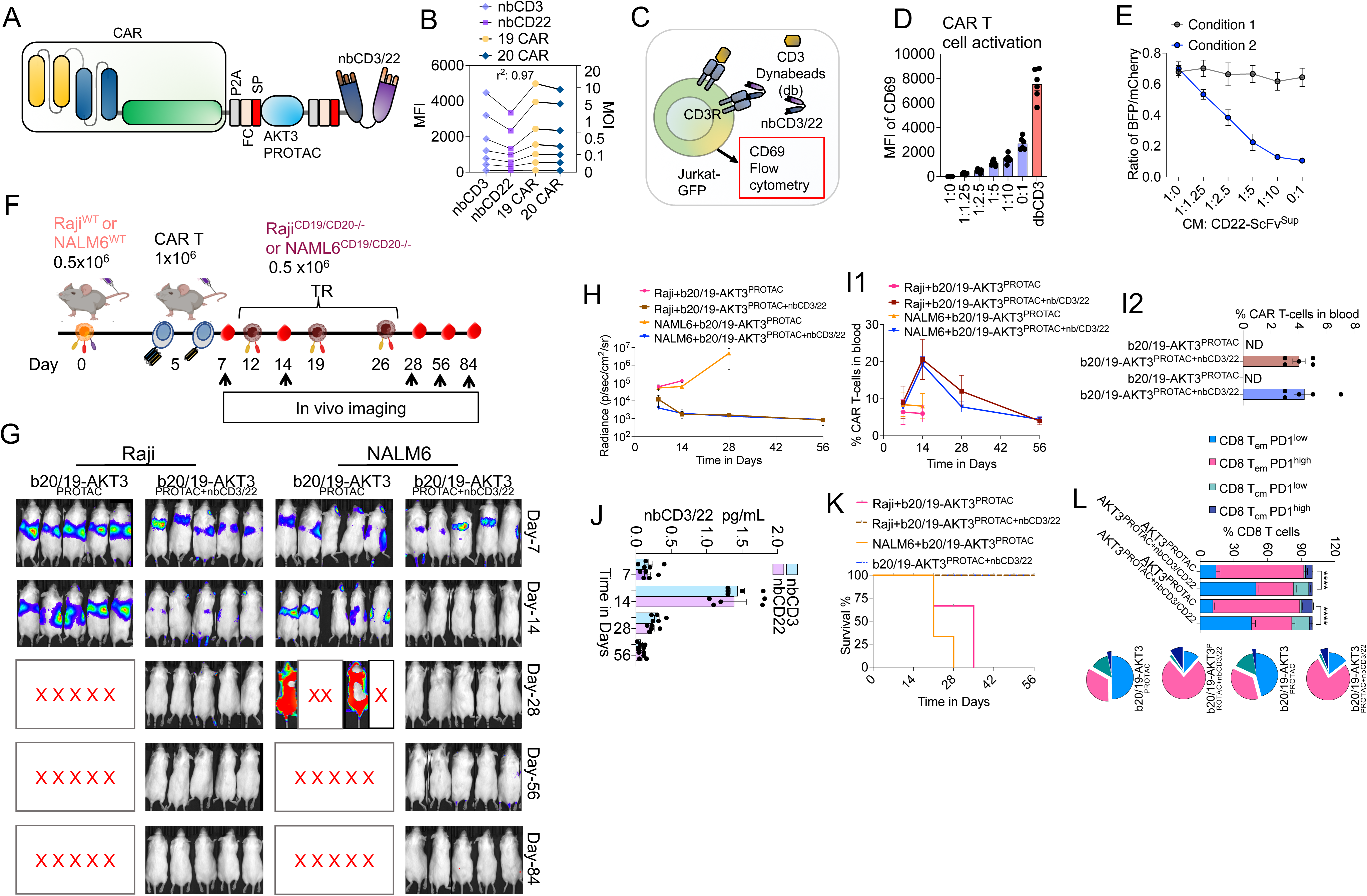
Trispecific CAR-T cells secreting CD3/CD22 targeted BITE enhance anti-tumor efficacy and persistence. (A) Schematic of the engineering strategy for trispecific CAR-T cells, integrating b20/19-AKT3^PROTAC^ with a secretory BiTE module consisting of nanobodies targeting CD3 and CD22 (nbCD3/22). (B) Correlation of expression of nbCD3, nb22, CD19 CAR, and CD20 CAR at various MOIs. The cells were treated with Brefeldin and data was obtained using intracellular flow cytometry. (C) Experimental setup for T cell activation, using Jurkat-GFP cells and Dynabeads (db) coated with CD3 to assess secreted nbCD3/22 functionality via flow cytometry. (D) Dose-dependent T cell activation (CD69 expression) in response to culture supernatants with nbCD3/22, using db coated with CD3 for validation. (E) HEK-293T synNotch reporter assay shows dose-dependent inhibition of CD22-CAR signaling by nbCD22 in CAR-T cell supernatants, confirming BiTE functionality under two conditions. (F) Experimental timeline for in vivo CAR-T cell therapy study in Raji^WT^ or NALM6^WT^ model followed by CAR-T cell administration and TR with Raji^CD19/CD20-/-^ or NALM6^CD19/CD20-/-^ cells (G) Bioluminescence imaging of Raji and NALM6 tumor-bearing mice treated with b20/19-AKT3^PROTAC^ or b20/19-AKT3^PROTAC+nbCD3/22^ CAR-T cells, monitored from Day 7 to Day 84. (H) Quantified tumor radiance over time, showing sustained tumor control in Raji and NALM6 models with b20/19-AKT3^PROTAC+nbCD3/22^. (I1) Percentage of CAR-T cells in the blood of Raji and NALM6 tumor-bearing mice treated with b20/19-AKT3^PROTAC^ or b20/19-AKT3^PROTAC+nbCD3/22^, measured over 56 days (I2) Bar graph of CAR-T cell populations in blood at various time points. (J) Levels of nbCD3/22 (pg/mL) in the blood of Raji and NALM6 tumor-bearing mice treated with b20/19-AKT3^PROTAC+nbCD3/22^, measured over 56 days, showing sustained secretion. (K) Kaplan-Meier survival curves demonstrating improved survival with nbCD3/22-modified CAR-T cells. (L) Bar graph and pie charts compare b20/19-AKT3^PROTAC^ and b20/19-AKT3^PROTAC+nbCD3/22^, showing various memory T cell subsets over time (n=5) in all conditions. Data represents mean ± SEM. ****p < 0.001. A non-parametric t-test was used for statistical analysis between groups.

To determine whether the secretory nbCD3 and nbCD22 are functional, we made a T cell activation model to study nbCD3 (**Figure 7C**) and HEK-293T reporter system based on synNotch system developed previously^39^ (**Figure S15**). We then tested the supernatant obtained from CAR-T cells expressing the b20/19AKT3^PROTAC/nbCD3/22^. A dose-dependent increase in CAR-T cell activation was observed with nbCD3, which was also confirmed by Dynabeads (db) for CD3 (**Figure 7D**). Similarly, the reporter system showed a dose dependent suppression of the CD22 CAR mediated signaling in HEK-293T cells, which was neutralized by the supernatant containing nbCD22 (**Figure 7E)**. These results proved the functional efficiency of the b20/19AKT3^PROTAC^ CAR-T cells expressing nbCD3/22.

Next, we assessed the preclinical efficacy of the nbCD3/CD22 BITE-secreting CAR-T cells using the TR model with Raji or NALM6 cells engineered to lack CD19 and CD20 but retain CD22 expression (**Figure 7F**). Mice treated with b20/19^AKT3PROTAC+nbCD3/22^ CAR-T cells exhibited robust and sustained anti-tumor responses over time (**Figure 7G, H**). This was accompanied by enhanced CAR-T cell persistence (**Figure 7I**). Importantly, the nbCD3/CD22 BITE was detectable in circulation alongside CAR-T cells (**Figure 7J**). Following tumor rechallenge, levels of both BITE and CAR-T cells increased, and subsequently declined in parallel, suggesting coordinated regulation of persistence and effector function (**Figure 7I, J**). These molecular effects translated into significantly improved survival in mice treated with b20/19^AKT3PROTAC+nbCD3/22^ CAR-T cells (**Figure 7K**). This enhanced therapeutic outcome was further supported by an increased frequency of memory CAR-T cell subsets exhibiting reduced exhaustion marker expression (**Figure 7L**). Importantly, toxicity assessment of the trispecific CAR-T cells revealed no significant adverse effects, and cellular persistence remained intact (**Figure S16, S17**).

## Discussion

A major challenge in achieving durable responses with CAR-T cell therapy is limited persistence, particularly in multi-antigen constructs such as bispecific and trispecific CARs. While considerable efforts have focused on optimizing co-stimulatory domains, the structural behavior of these complex designs remains insufficiently characterized. Elements such as multiple antigen-binding domains like single-chain fragment variables (scFvs), glycine-serine linkers, and modular co-stimulatory motifs can significantly influence CAR stability, surface expression, and function, ultimately impacting therapeutic efficacy. We have addressed these issues by systematically screening a library of mono, bi, and trispecific CAR constructs, varying antigen specificities, co-stimulatory domains (CD28, 4-1BB, ICOS), and linker lengths. We developed an integrated AI-driven pipeline, CARMSeD that predicts functional liabilities, including self-activation signaling potential, exhaustion signatures and CAR-T cell death. Unlike previous AI models, which focus on single readouts such as activation^40,41^ or individual domain properties like co-stimulation^42^, our platform integrates multiple structural parameters and functional outputs to enable a more comprehensive prediction of CAR behavior and CAR-T cell function. Additionally, we developed a computational model to estimate ScFv-ScFv interaction propensities, addressing a critical yet understudied determinant of ligand-independent signaling in multiple car designs. Based on this extensive in silico screening, we identified CAR designs with reduced self-activation signaling and improved functional properties. Among top candidates, a bispecific CD20/19 CAR consistently showed high expression, low CARMSeD scores, and strong in-vitro performance across proliferation, cytokine secretion, and cytotoxicity assays. These findings highlight the critical role of rational structural design in improving CAR-T cell efficacy and persistence, consistent with previous studies linking CAR architecture to therapeutic outcomes^43,44^.

To further improve the durability of these multi-antigen CAR-T cells, we focused on the pathways that are integral to the T cell memory. Based on the RNA Seq analysis, AKT3 showed differential expression during CAR-T cell exhaustion. As previous studies have shown AKT signaling critical for the T cell memory formation, T cell exhaustion, and metabolic fitness we explored the specific role of AKT3 further^33–35^. To functionally interrogate role of AKT3, we have employed a PROTAC-based strategy for selective degradation of AKT3. PROTACs represent a novel class of molecules that induce the targeted degradation of specific proteins via the ubiquitin-proteasome pathway^45,46^. Unlike traditional inhibitors that merely block protein function, PROTACs hijack E3 ligases to ubiquitinate the target protein, ensuring complete and irreversible removal^45,47^. In our study, PROTAC-mediated degradation of AKT3 led to a marked enhancement in CAR-T cell persistence, metabolic reprogramming, and resistance to exhaustion. Mechanistically, AKT3 degradation promoted a shift in T cell metabolism from glycolysis to OXPHOS, a hallmark of long-lived T_cm_ and T_scm_ subsets^48–50^. These subsets are essential for sustained antitumor surveillance and re-expansion upon antigen re-encounter. Our data revealed a marked enrichment of T_cm_ and T_em_ subsets within AKT3-targeted CAR-T cell populations, which strongly correlated with sustained cytotoxic activity and extended survival in murine models. These memory-enriched CAR-T cells demonstrated enhanced metabolic fitness, which translated into markedly improved tumor control across multiple B-cell leukemia and lymphoma models. Importantly, this enhanced persistence, and functionality aligns with emerging evidence that PROTACs, particularly bioPROTACs, can be harnessed to precisely degrade intracellular signaling components such as ZAP70, enabling fine-tuned modulation of CAR signaling^51^. This approach offers a novel, programmable strategy to enhance therapeutic efficacy while improving safety by limiting off-target or excessive activation. The most prominent findings in our study was the upregulation of FOXO4 following AKT3 degradation. FOXO4 is a transcription factor involved in autophagy, metabolic homeostasis, and longevity, and plays a key role in downregulating mTOR signaling^52,53^. Given our previous findings that mTOR suppression is critical for CAR-T cell memory formation, we were prompted to investigate the functional contribution of FOXO4 in this context more closely^32^. Unlike its family members FOXO1 and FOXO3, which are well-established regulators of T cell memory formation^37,54,55^, FOXO4 has not been extensively studied in the context of T cells. However, our findings reveal that FOXO4 plays a pivotal role in maintaining CAR-T cell fitness. We observed that AKT3 suppresses FOXO4 activity through phosphorylation. Targeted degradation of AKT3 led to increased FOXO4 expression, which was associated with enhanced memory T cell formation, improved mitochondrial function, and reduced exhaustion. Importantly, the knockdown of FOXO4 abolished the functional benefits conferred by AKT3 degradation, confirming FOXO4 as a central mediator in this regulatory axis.

We further translated this strategy into a trispecific CAR-T platform targeting CD19, CD20, and CD22. This platform included a secretory nanobody-based on CD3/CD22 T cell engager to address dual antigen escape, an emerging mechanism of relapse in B-cell malignancies, besides CD19 negative relapse^56^. When combined with AKT3-targeted PROTAC treatment, these trispecific CAR-T cells exhibited sustained expansion, superior tumor control, and long-term persistence even in antigen-heterogeneous settings. This triple-modality design proved effective in overcoming both structural and functional limitations inherent to complex CAR formats.

Importantly, the benefits of the AKT3-FOXO4 axis were not limited to hematological cancers. We extended our investigation to solid tumors, where CAR-T cell therapy has historically been less effective due to hostile microenvironments and poor persistence^57,58^. CAR-T cells targeting Claudin18.2 (for gastric cancer) and EGFR (for NSCLC) were modified with AKT3-PROTAC treatment. These cells showed improved trafficking, reduced exhaustion markers, and superior tumor regression in xenograft models, suggesting that this approach has broad applicability across cancer types.

Beyond AKT3, the PROTAC platform offers the flexibility to target other regulators of CAR-T cell exhaustion and immune evasion. For example, PROTACs directed against PD-1, TIGIT, or LAG-3 may enable real-time modulation of inhibitory checkpoint receptors, thereby enhancing T cell activity without the need for systemic checkpoint blockade^59,60^. Moreover, transient PROTAC exposure can fine-tune CAR-T cell phenotype without permanently altering the genome, offering a safer alternative to CRISPR or other gene-editing tools. Our findings position PROTAC technology as a versatile and powerful tool for next-generation CAR-T cell engineering. When integrated with rational CAR design, co-stimulatory optimization, and antigen multiplexing, PROTAC-mediated protein degradation and recently developed synthetic gene switches holds the potential to surmount long-standing limitations of CAR-T therapy^61^. These include not only antigen escape and exhaustion but also poor metabolic fitness and short-lived memory responses^32,62^.

In conclusion, our study introduces a multifaceted strategy combining high-throughput CAR screening, AI-assisted design, and PROTAC-based intracellular signaling modulation to generate functionally superior CAR-T cells. Targeting AKT3 via PROTACs significantly improved memory differentiation, metabolic robustness, and long-term persistence, which are crucial determinants of therapeutic success of multi-antigen directed CAR-T cells. The concurrent upregulation of FOXO4 further highlights a novel regulatory axis in T cell biology. This integrative approach represents a critical step forward in addressing the twin challenges of antigen escape and CAR-T cell exhaustion and has the potential to extend CAR-T efficacy across both hematologic and solid malignancies.

## Methods

### Cell line details

The cell lines K562, HEK-293T, Raji, NALM6, Jurkat-GFP, AGS, A549 and Daudi were purchased from the American Type Culture Collection (ATCC) and cultured following ATCC-recommended protocols. HEK-293T, AGS cells were maintained in DMEM with 10% FBS at 37°C, 5% CO₂, and 95% humidity, with sub culturing at 70–80% confluency using trypsin. Raji, K562, Jurkat, NALM6, and Daudi cells were cultured in RPMI-1640 medium with 10% FBS under similar conditions, with Raji and Daudi cells sub-cultured at 2–3 × 10^6^ cells/mL and Jurkat, K562 and NALM6 at 1–2 × 10^6^ cells/mL. Similarly, A549 cells were cultured under same conditions in F12/K media with 10% FBS.

### Human Samples

Peripheral blood mononuclear cells (PBMCs) were obtained from healthy adult donors between 20 to 55 years of age, with 20 male and 23 female samples obtained at Apollo Indraprastha Hospital, New Delhi, in accordance with protocols approved by the Institutional Review Board. Additionally, PBMCs were collected from 5 patients, following written informed consent (as described in our previous article^32^) and adherence to the hospital’s ethical and regulatory standards. The blood samples utilized were surplus specimens that would otherwise have been discarded after marker analysis.

### Integrated Meta-Analysis of CAR-T Therapy Resistance Mechanisms

A meta-analysis was conducted to assess relapse patterns and structural vulnerabilities in CAR-T cell therapy, specifically targeting CD19 and CD20 antigens in mono and bispecific constructs. Existing clinical data from 4,129 patients treated with CAR-T cells were compiled from peer-reviewed publications indexed on PubMed, registered clinical trial databases (including studies from the USA, China, and Europe), and preprint servers such as bioRxiv (**Table 1**). Patient-level and cohort-level information were extracted to identify relapse rates, with a specific focus on CD19-negative tumor recurrences. Studies involving both monospecific and multi-antigen (bi and trispecific) CAR-T therapies were included. The compiled data were analyzed to evaluate the frequency of antigen escape and CAR-T cell persistence.

### Profiling of CD19, CD20 and CD22 receptor expression in ALL patients

Flow cytometry data from ALL patients were collected from Sir Ganga Ram Hospital (Delhi), Medanta Hospital (Gurugram), and Shalby Sanar International Hospital (Gurugram), India, following ethical approval and informed consent from all patients. Samples from over 129 ALL patients were included to assess surface expression of CD19, CD20 and CD22 antigens (**Table 4**). The data were analyzed data were categorized based on intensity and distribution into four groups: positive (strong, uniform expression), dim (low-intensity expression), negative (no detectable expression), and heterogeneous (variable expression within the cell population).

### Animal Experiments

NOD.Cg-Prkdc scid Il2rg tm1Wjl /SzJ (NSG) and NOD-Prkdcem26Cd52Il2rgem26Cd22/NjuCrl (NCG), aged 6–8 weeks, were purchased from The Jackson Laboratory and Charles River Laboratories, respectively and housed in the Laboratory. Additionally, 4–6-week-old female SCID-beige mice (strain C.B-Igh-1b/GbmsTac-Prkdcscid-Lystbg N7) were sourced from Taconic Biosciences for CRS studies. These animals were also kept under a 12 hrs light/dark cycle at 22°C with free access to food and water. All animal experiments were conducted following approval from the respective Ethics Committee for Animal Experiments. All animal experiments were approved by Institutional Animal Ethics Committee; CSIR-Institute of Chemical Biology, Kolkata, West Bengal, India; Regional Centre for Biotechnology, Faridabad, Haryana, India; and Adgyl Lifesciences, Hyderabad, India.

### Reporter Cell lines and peptide screening

Reporter systems were established as follows: A GFP-AKT3-mCherry reporter construct was generated in HEK293T cells via transduction with a lentiviral vector encoding mCherry-AKT3. The mCherry-AKT3 fusion plasmid was assembled using synthetic gene fragments (Genscript) and cloned using Gibson Assembly. The lentiviral vector used for transduction was prepared as previously described^32^. Stable GFP-AKT3-mCherry-expressing cell lines were established through single-cell sorting using FACS. Peptides targeting AKT3 were tested in these cells. Transfections were performed in 12-well plates at a density of 2×10^5^ cells per well using 1 µg of plasmid DNA and Lipofectamine 3000 (Thermo Fisher Scientific). Cells were treated with PROTAC peptides (0.1–10 µM) for 24 hours, followed by assessment of AKT3 degradation using confocal microscopy (Zeiss LSM 880) and western blotting with anti-AKT3 (Cell Signaling Technology, #14982) and anti-GAPDH (Sigma-Aldrich, G8795) antibodies.

A SynNotch reporter system was developed using SynNotch plasmids obtained from Addgene (Plasmid #79125 and #79130), as described previously^39^. Synthetic gene fragments encoding the CD22 receptor were synthesized and transduced into HEK293T-GFP cells to create sender cells. A stable clone (CD22R-HEK293T-GFP) was selected by FACS. A separate synthetic fragment encoding CD22-scFv was cloned into the SynNotch vector (Plasmid #79125), and the resulting lentiviral construct was used to transduce HEK293T cells. Additionally, a vector encoding pHR_Gal4UAS_tBFP_PGK_mCherry (Plasmid #79130), enabling inducible BFP expression and constitutive mCherry expression, was used to generate receiver cells. Stable receiver HEK293T cells co-expressing both plasmids were isolated by FACS.

For the co-culture assay, sender and receiver cells were plated at a 1:1 ratio and treated with b20/19-AKT3^PROTAC^ ^+ nbCD3/22^ supernatants (dilutions ranging from 1:2 to 1:10) for 48 hours. BFP expression was assessed via confocal microscopy to confirm nbCD22-mediated inhibition of CD22-CAR signaling.

A Jurkat-GFP reporter assay was developed to assess nbCD3 functionality. Jurkat-GFP cells (1×10^5^ per well) were plated in 96-well plates and incubated with the same supernatants (1:2 to 1:10 dilutions) for 24 hours. Dynabeads CD3/CD28 (Thermo Fisher Scientific) served as positive controls. Cells were stained for the T cell activation marker CD69 using an APC-conjugated anti-CD69 antibody (BD Biosciences, #555533). Flow cytometry was used to measure CD69 expression, confirming T cell activation. Further, to model CD20 heterogeneity, K562 cells were transduced with lentiviral vectors encoding CD20 at varying expression levels (low, medium, high), sorted by FACS based on CD20 expression (anti-CD20-PE, BD Biosciences).

### CAR Library Construction and in-vitro screening

A library of 10,824 chimeric antigen receptor (CAR) constructs was systematically generated, encompassing mono, bi, and trispecific formats targeting CD19, CD20, and CD22 antigens. Constructs were designed with variability in glycine-serine (G_4_S) linker lengths (3, 4, or 5 G_4_S repeats connecting scFvs and VH/VL domains), co-stimulatory domains (CD28, 4-1BB, or ICOS), transmembrane domains (predominantly ICOS-derived), and a CD3ζ signaling domain for T cell activation. From this library, 1,452 constructs were selected for in-vitro functional screening based on diversity in structural configurations. Primary human T cells were isolated from peripheral blood mononuclear cells (PBMCs) of three healthy donors using magnetic bead separation with CD3/CD28 Dynabeads (Thermo Fisher Scientific). T cells were activated with anti-CD3/CD28 antibodies (Thermo Fisher Scientific) at 1 µg/mL for 48 hours and transduced with lentiviral vectors encoding the selected CAR constructs at a multiplicity of infection (MOI) of 5, in the presence of polybrene (8 µg/mL, Sigma-Aldrich). Transduced T cells were expanded in RPMI-1640 medium (Gibco) supplemented with 10% fetal bovine serum (FBS, Gibco) and 100 IU/mL recombinant human IL-2 (PeproTech) for 7 days. Functional screening evaluated three parameters: T cell activation (signal-1) via CD69 expression, exhaustion (signal-2) via PD-1 expression, and apoptosis (signal-3) via 7-AAD staining, all measured by flow cytometry using anti-CD69-FITC, anti-PD-1-PE (BD Biosciences), and an 7-AAD apoptosis detection kit (Thermo Fisher Scientific). Signal intensities were categorized into low (L), medium (M), and high (H) levels based on cutoffs established using a second-generation monospecific CD19 CAR as a reference. A scoring system was applied to classify CARs into low, medium, and high CAR-mediated self-destruction (CARMSeD) categories, reflecting their propensity for self-activation and dysfunction

### AI-Guided Screening with CARMSeD Model

To facilitate downstream classification, signal values were discretized into categorical bins: Signal-1 was divided into low, medium, and high using thresholds at 0.5 and 1.0; Signal-2 and Signal-3 were binned using thresholds at 0.25 and 0.5. Each CAR sequence was then assigned to one of three functional synergy categories, L-CARMSeD, M-CARMSeD, or H-CARMSeD based on a rule-based mapping of all valid signal triplets. This labeling scheme was designed to reflect combinatorial synergy and activation potency. CAR sequences were tokenized using a fixed amino acid vocabulary of 20 standard residues, with unknown or non-canonical residues assigned to a special out-of-vocabulary token. All sequences were truncated or zero-padded to a uniform length of 1,024 residues. The model, termed CARMSeD, was implemented in TensorFlow and structured as a multi-task deep neural network. The architecture began with a trainable embedding layer (dimensionality 128), followed by two one-dimensional convolutional layers, each with 256 filters and a kernel size of 5, using ReLU activations. A max-pooling operation was followed by a global max pooling layer to condense spatial representations. The resulting latent feature map was branched into two task-specific heads: a classification head with SoftMax activation to predict the CARMSeD class, and a regression head with linear activation to predict the continuous values of Signal-1, Signal-2, and Signal-3.

Model training was conducted for up to 50 epochs using the Adam optimizer with a batch size of 64. An early stopping criterion based on validation classification accuracy was employed to prevent overfitting. The composite loss function combined categorical cross-entropy (for classification) and mean squared error (for regression), with loss weighting of 3:1 to prioritize accurate functional classification. All signal values were normalized to the [0, 1] range using MinMaxScaler prior to training. The dataset was split into training and validation cohorts in an 80:20 ratio using stratification based on the CARMSeD class and discretized Signal-3 values to preserve class balance. Model performance was evaluated using classification accuracy, confusion matrices, mean squared error, and the coefficient of determination (R²) for regression tasks. After training, the CARMSeD model was applied to held-out test datasets to generate predictions for both categorical class labels and signal values. Model outputs were visualized using Python libraries matplotlib and seaborn. Visual inspection included measured vs. predicted scatter plots for each signal axis, residual error distributions, and class-specific confusion matrices to characterize error structure and functional resolution across the prediction space.

### Primary cell isolation and transduction

The PBMCs were extracted from whole blood samples by density gradient centrifugation using Ficoll-Paque (Cytiva) as per the manufacturer’s guidelines. Subsequently, CD4/CD8 magnetic beads were used to isolate T-cells. Similarly, CD4 and CD8 magnetic beads (Miltenyi Biotec) were used to isolate pure CD4^+^ and CD8^+^ T-cell populations. These T-cells were then activated using anti-CD3/CD28 stimulation (T-cell TransAct, Miltenyi Biotec) and cultured in TexMACS T-cell culture medium supplemented with 100 IU/mL of human recombinant IL-7 and IL-15 (Miltenyi Biotec). Following 24 hrs of activation, the T-cells were transduced with lentiviral concentrate at MOI of 5 and incubated for an additional 48 hrs before media change. Fresh medium was replenished every 2-days throughout the experiment. The percentage of transduced T-cells was determined using the CD19 CAR detection reagent (Miltenyi Biotec) following the manufacturer’s instructions, and flow cytometry analysis was done after day 5 of transduction, unless otherwise specified.

Patient blood samples were processed to isolate PBMCs using Ficoll density gradient centrifugation. PBMCs were further separated into T-cells and B-cells. For T-cell isolation, standard procedures were followed, while for B-cell isolation, magnetic beads conjugated to CD19 antibodies (Miltenyi Biotech) were utilized. The isolated B-cells were confirmed via flow cytometry using CD19-specific antibodies. After isolation, the cells were cultured in RPMI medium supplemented with 10% FBS and subsequently co-cultured with CAR-T-cells, which were generated from the patient-derived T-cells. All procedures adhered strictly to ethical guidelines and regulatory standards to ensure patient safety and data integrity.

### Humanization and computational validation of scFvs

To evaluate the structural and dynamic properties of bispecific (anti-CD19/CD20) and trispecific (anti-CD19/CD20/CD22) CAR constructs, a computational pipeline was established. Initial murine scFv sequences and structures for anti-CD19 and anti-CD20 were retrieved from the Protein Data Bank (PDB IDs: 7URV and 6VJA, respectively), while the humanized anti-CD22 scFv sequence and structure were obtained from PDB ID: 7O52. All possible bispecific and trispecific constructs were generated using AlphaFold2 (DeepMind) for ab initio structure prediction, followed by structural refinement and visualization in UCSF Chimera (v1.16). Molecular dynamics (MD) simulations were performed to investigate intramolecular interactions between scFvs within these constructs using Amber22 with the AMBER ff14SB force field and TIP3P water model. The simulation protocol consisted of three stages: energy minimization, heating, and production. Systems were minimized over 10,000 steps, with 2,000 steps of steepest descent followed by 8,000 steps of conjugate gradient. The minimized systems were heated to 300 K over 150 ps under NVT conditions, followed by a 200 ns production run under NPT conditions at 1 bar and 300 K, using a 2 fs timestep. Trajectory analysis was conducted using CPPTRAJ (AmberTools) to calculate root mean square deviation (RMSD) and intramolecular hydrogen bond interactions, focusing on complementarity-determining regions (CDRs).

Humanization of the murine anti-CD19, and anti-CD20 scFvs was performed using BioPhi (v1.0), a web-based tool offering two approaches: CDR grafting and the Sapiens iterative humanization method. Both CDR grafting and Sapiens method were used to generate library of humanized sequences. From those, the one with high humanness in both approaches were selected to iteratively optimize framework regions while preserving CDR integrity, generating humanized scFv variants with high sequence identity to human germline frameworks. The resulting humanized scFvs were modeled using the corresponding murine structures (PDB IDs: 7URV, and 6VJA) as templates in MODELLER (v10.1), producing 10 homology models per scFv, with the best model selected based on the DOPE score. These models were docked against their respective antigens (CD19, CD20) using the HADDOCK webserver (v2.4), employing default parameters to predict/generate best fitting complexes. To compare the binding affinities of murine and humanized scFv variants, MD simulations were conducted using the same AMBER ff14SB force field and TIP3P water model. Systems were prepared as described above, but production runs were extended to 300 ns to ensure convergence of binding interactions. Trajectory analysis included RMSD, hydrogen bond analysis, per-residue decomposition, and enthalpy calculations using the MM-GBSA method in AmberTools23, validating structural stability, interaction profiles, and binding affinity of the humanized variants.

The bispecific and trispecific combination were then modelled with optimized humanized scFvs via homology modelling on previously described structures, to preserve spatial orientation between scFvs in the bispecific construct. The resulting bispecific structure was subjected to MD simulation following the same protocol as above, with a 200 ns production run. Final trajectory analysis focused on RMSD and intramolecular hydrogen bond interactions within CDRs to confirm structural stability and inter-scFv interactions critical for antigen recognition.

### In-vitro functional validation

K562 cells were engineered to stably express CD19, CD20, and CD22 individually, as well as in combination, using lentiviral transduction. Synthetic gene fragments encoding CD19 (NM_001385732.1), CD20 (NM_021950.4), and CD22 (NM_001185099.2) were cloned into a previously described lentiviral backbone^32^. Stable cell lines were established by FACS sorting using anti-CD19, anti-CD20, and anti-CD22 antibodies, and expression of each antigen was confirmed by flow cytometry. Cytotoxicity assays were conducted by co-culturing CAR-T cells (b20/19, b22/19, and t20/19/22) with K562 target cells at effector-to-target (E:T) ratios of 1:1, 2.5:1, 5:1, and 10:1 for 24 hours. Tumor cell lysis was quantified using 7-AAD staining. T cell proliferation was assessed as previously described^32^. Raji cell knockout clones were generated using gene-specific gRNAs targeting CD19, CD20, and CD22. A detailed protocol for the generation of these stable Raji knockout cell lines is currently under review in a separate manuscript.

### CAR-T-cell proliferation, persistence and cell death

CAR-T cell proliferation was assessed using the CellTrace™ CFSE Cell Proliferation Kit (Thermo Fisher Scientific). Briefly, CAR-T cells were stained with CFSE dye, and proliferation was monitored at various time points (Days 3, 5, 7, 9, 11, 13, 15, 17, and 19), as previously described^32^. Initially, CAR-T cells were harvested and washed to ensure viability and purity. CAR-T cell persistence was measured by counting cell numbers on Day 0 (prior to co-culture) and on Days 7, 13, 21, and 28 following co-culture with Raji cells. Co-cultures were performed on alternate days (Days 6, 8, 10, 12, 14, 16, 18, and 20). The assay methodology followed our previously published protocol^32^.

Cell death analysis was conducted using 7-AAD staining, followed by acquisition on a flow cytometer, as previously described^32^. Data were presented as either representative flow cytometry histograms showing the percentage of mean fluorescence intensity of 7-AAD, or as bar graphs indicating percent cell death.

### Transcriptomic and functional analysis of AKT3

CAR-T cells were isolated from control and tumor-rechallenged (TR) mice (n=5 per group) treated with b20/19 CAR-T cells. RNA was extracted using QIAGEN Pure mRNA beads, following further enrichment and heat fragmentation, RNA was reverse-transcribed into cDNA, employing RNase H-Reverse Transcriptase and primers for strand-specific ligation. and transcriptomic profiling was performed via RNA sequencing on an Illumina NovaSeqTM 6000. Differential gene expressions were analyzed using iGeak_RNA-seq, focusing on pathways such as oxidative phosphorylation, FOXO signaling, and autophagy. The RAN Seq analysis was done as described by us previously^32^. AKT3 upregulation was confirmed by flow cytometry using anti-AKT1/2/3-FITC, ThermoFisher, #BS-6951R-FITC. AKT3 overexpression was achieved by co-transducing CAR-T cells with a lentiviral vector encoding AKT3 via pLenti-AKT3, while knockdown was performed using shRNA **(Table 5 & Table 6)**.

### PROTAC design and validation

AKT3-targeting PROTAC peptides were designed using RF diffusion modeling with AlphaFold (https://github.com/RosettaCommons/RFdiffusion), yielding a library of peptides of which top five candidate peptides (AKT3_P1 to AKT3_P5) and and a non-targeting peptide (NTP) as a control were selected based on the sequence diversity, secondary structure and complex structure confidence score. Each peptide was fused to a TAT-derived cell-penetrating peptide (CPP) and a VHL E3 ligase ligand via a flexible linker to facilitate intracellular delivery and ubiquitin-mediated degradation. Degradation efficacy was evaluated in HEK293T cells (ATCC, CRL-3216) stably expressing a GFP-AKT3-mCherry reporter, seeded at 2×10⁵ cells per well in 12-well plates. Cells were treated with peptides at concentrations ranging from 0.1 to 10 µM for 24 hours in DMEM (Gibco) supplemented with 10% FBS. The peptides were synthesis from GenScript at a concentration of 2 to 4mg each. AKT3 degradation was quantified by fluorescence microscopy on a Zeiss LSM 880 confocal microscope, measuring the mCherry/GFP fluorescence ratio with ImageJ (v1.53), and by western blotting using antibodies against AKT3 (1:1000, Cell Signaling Technology, #14982), GFP (1:2000, Abcam, ab290), and GAPDH (1:5000, Sigma-Aldrich, G8795) as a loading control. The top-performing peptide, P2, was initially tested in GFP expressing CAR-T cells and endogenous AKT3 was detected using rabbit anti-AKT3 primary antibody followed by staining with Alexafluor 543-conjugated secondary anti-rabbit antibody. Notably, AKT3-PROTAC (without CPP) was integrated into the b20/19 CAR construct downstream of the CAR sequence using a P2A self-cleaving peptide linker, cloned into a pLenti vector. Primary T cells transduced with this construct were co-cultured with CD19^-/-^ Raji cells at an effector-to-target (E:T) ratio of 2.5:1 for 24 hours, and AKT3 degradation was confirmed by fluorescence microscopy and western blotting. Cytotoxicity was assessed against wild-type Raji cells and patient-derived leukemia samples (n=5) at the same E:T ratio for 24 hours, with target cell survival quantified by flow cytometry using 7-AAD staining (Thermo Fisher Scientific). CAR-T cell persistence was monitored over 15 days in a tumor rechallenge model with CD19-/-Raji cells, while T cell subsets (T_cm_, T_em_) and exhaustion markers (PD-1) were analyzed by flow cytometry using anti-CD62L-FITC, anti-CD45RO-PE, and anti-PD-1-APC (BD Biosciences). Metabolic profiles, including extracellular acidification rate (ECAR) and oxygen consumption rate (OCR), were measured as described by us previously^32^.

### Functional and biochemical assays

Immunoblotting for AKT3 and GAPDH was performed as previously described^32^. Similarly, RT-qPCR assays were conducted following our established protocols. Primers used for AKT3 and FOXO4 are listed in **Table 5**. Total RNA was extracted using the RNeasy Mini Kit (Qiagen) according to the manufacturer’s instructions, and RNA integrity was assessed using a NanoDrop 2000 spectrophotometer (Thermo Fisher Scientific). FOXO4 expression was quantified by quantitative real-time PCR (qRT-PCR) using a QuantStudio 3 system (Applied Biosystems), with primers designed via Primer-BLAST (**Table 5**).

FOXO4 and AKT3 knockdown was performed using pLKO.1 lentiviral vector carrying FOXO4-shRNA and AKT3-shRNA, with efficiency confirmed by qRT-PCR and western blotting as described by us previously.

### CAR Binding Analysis

Binding affinity analysis of CAR ScFvs targeting CD19, CD20, and CD22 was performed using a flow cytometry-based method (BD FACSLyric). T cells were transduced with lentiviral vectors encoding the respective CAR ScFvs. Following transduction, cells were fixed and stained using anti-CAR antibodies that recognize the extracellular domain (ECD) of the respective receptors: CD19 (Cat No. 130-129-550) and CD22 (Miltenyi Biotec). The CAR ECDs were biotin-conjugated, and detection was carried out using PE-conjugated anti-biotin REAfinity antibody (clone REA746). For CD20 binding analysis, a plasmid encoding the CD20 extracellular domain (ECD) was generated. A synthetic CD20 ECD fragment was cloned into the vector pD649-HAsp-CD22-Fc (DAPA)-AviTag-6xHis (Addgene Plasmid #156940). This construct was used to purify recombinant CD20 protein conjugated to biotin. The CD20-biotin protein was then used for assessing binding to CD20-targeting CAR ScFvs. Protein purification, expression and isolation of recombinant CD20 were performed according to our previously published protocol^63^. Binding affinity data were analyzed using FlowJo software and presented as percentage binding.

### Flow Cytometry Analysis

Flow cytometry was performed using BD FACS Aria, Accuri, Lyric systems, and Beckman coulter CytoFLEX Flow Cytometer and data were analyzed with FlowJo or CytExpert software. The flow cytometry was performed as described by us previously^32^. Briefly, CD19, CD22 CAR expression was evaluated using CD19 and CD20 CAR detection antibodies and CD22 CAR expression (Miltenyi Biotec) was evaluted using Protein L-APC (Cell signaling) followed by PE-conjugated anti-biotin secondary antibodies (Miltenyi Biotec). As described previously, T cell immunophenotyping and functional marker analyses (e.g., Granzyme B, PD-1, LAG-3) were performed using fluorophore-conjugated antibodies. Antibodies against CD45RO, CCR7, and PD-1 to identify memory and exhausted T cell subsets. Cell death was determined using 7-AAD staining. For FOXO4 and phospho-FOXO4 (Ser193) levels, cells were fixed/permeabilized with the Foxp3/Transcription Factor Staining Buffer Set (eBioscience) and stained with anti-FOXO4-APC (Abcam, ab211689) and anti-phospho-FOXO4-PE. The data was analysed using FlowJo software and presented as mean fluoresce intensity (MFI) unless specified.

### Super-Resolution Microscopy

Super-resolution microscopy was done to visualize low-level CD20 expression in K562 cells. A total of 1.5 × 10^6^ cells were harvested from actively growing cultures and washed twice with 1X phosphate-buffered saline (PBS) (Thermo Fisher Scientific, Cat# 10010023). Cells were then incubated with a fluorophore-conjugated CD20 antibody (Sigma-Aldrich, Cat# SA134700118) prepared at a dilution of 2 µL antibody per 100 µL 1× PBS, following the manufacturer’s instructions. The staining was carried out at 4°C for 20 minutes. Following staining, cells were washed twice with 1X PBS and subsequently fixed with 4% paraformaldehyde (PFA) at room temperature for 15 minutes. Fixed cells were washed and mounted onto poly-L-lysine–coated glass slides (Sigma-Aldrich, Cat# P0425-72EA) using ProLong™ Gold Antifade Mountant (Thermo Fisher Scientific, Cat# P36971). The slides were allowed to dry overnight at room temperature in the dark. Imaging was performed using a Zeiss ELYRA PS.1 3D PALM, equipped with a 63X oil immersion objective (Plan-Apochromat 63×/1.40 Oil DIC; Filter: MBS-561 + EF BP 570–650/LP750; 100 mW laser). Structured Illumination Microscopy (SIM) was carried out using a 561 nm laser line. Image stacks were acquired and processed using Zeiss Zen software. Final image analysis, including intensity profiling and comparative visualization was performed using ImageJ.

### In Vivo Studies

NOD/SCID/IL2Rγ-null (NSG) mice (6–9 weeks old, n=5 or 6 per group) were injected intravenously with 1×10^6^ Raji or NALM-6 cells expressing luciferase on day 0, followed by 1×10^7^ CAR-T cells (b20/19, b20/19-AKT3^PROTAC^, or b20/19-AKT3^PROTAC+nbCD3/22^) on day 5, unless specified. Tumor rechallenge with 1×10^6^ CD19^-/-^ or CD19/CD20^-/-^ cells was performed on days 12, 19, and 26 and so on. Tumor burden was monitored by bioluminescence imaging using an IVIS Spectrum, PerkinElmer after intraperitoneal injection of D-luciferin as per the protocol previously described^32^. CAR-T cell persistence and nbCD3/22 levels in peripheral blood were quantified by flow cytometry and ELISA, respectively. Survival was analyzed using Kaplan-Meier curves with the Mantel-Cox test. For solid tumor studies, AGS (gastric cancer) and A549 (NSCLC) xenografts were established, and CAR-T cells targeting Claudin18.2 or EGFR were administered, with efficacy assessed as above.

### Trispecific CAR-T Cell Engineering

Trispecific CAR-T cells were engineered by integrating the b20/19-AKT3^PROTAC^ construct with a secretory CD3/CD22 bispecific T cell engager (BiTE, nbCD3/22). The nbCD3 and nbCD22 nanobodies were humanized via CDR grafting using BioPhi (v1.0) to minimize immunogenicity while preserving antigen-binding affinity. Humanized nanobody sequences were synthesized (GenScript) and cloned into a pLenti vector downstream of the b20/19-AKT3^PROTAC^ sequence, separated by a P2A self-cleaving peptide to ensure co-expression, using Gibson Assembly (New England Biolabs). Constructs were sequence-verified by Sanger sequencing. Primary human T cells, isolated as described above, were transduced with the lentiviral vector at multiplicities of infection (MOIs) ranging from 2 to 10 in the presence of polybrene (8 µg/mL, Sigma-Aldrich), and expanded in RPMI-1640 medium (Gibco) supplemented with 10% FBS and 100 IU/mL IL-2 (PeproTech) for 7 days. Expression of nbCD3/22 was quantified on day 7 post-transduction by fluorescent ELISA using a custom anti-CD3/anti-CD22 detection system with a fluorescence VICTOR Nivo Multimode Microplate Reader (Revvity). Intracellular expression was confirmed by flow cytometry using anti-CD3-FITC (1:100, BD Biosciences, #555332) and anti-CD22-PE (1:100, BD Biosciences, #562859) antibodies after treating cells with Brefeldin A (5 µg/mL, Sigma-Aldrich) for 4 hours to block secretion, followed by fixation and permeabilization. Functionality of the secreted nbCD3/22 BiTE was assessed in two systems: (1) a Jurkat-GFP activation model, where Jurkat-GFP cells (1×10^5^ cells/well in 96-well plates) were incubated with CAR-T cell supernatants (1:2 to 1:10 dilution) for 24 hours, and T cell activation was measured by CD69 expression using anti-CD69-APC (1:100, BD Biosciences, #555533) via flow cytometry, with Dynabeads Human T-Activator CD3/CD28 (Thermo Fisher Scientific) as a positive control; and (2) a HEK293T SynNotch reporter assay, where sender cells (expressing CD22, SynNotch, GAL4-UAS-BFP, and GFP) and receiver cells (expressing mCherry) were co-cultured at a 1:1 ratio with CAR-T cell supernatants (1:2 to 1:10 dilution) for 48 hours, and BFP expression was quantified by confocal microscopy to confirm nbCD22-mediated inhibition of CD22-SynNotch signaling (Leica EM UC7, Leica, Wetzlar, Germany) as described by us previously^64^.

### Quantification and Statistical Analysis

Statistical analyses were performed using GraphPad Prism version 8.0 (GraphPad Software). Group differences were evaluated using non-parametric t-tests. Bar charts and associated P-values were generated with Prism. For comparisons involving more than two groups, a non-parametric one-way ANOVA was applied, while survival analyses between two groups were performed using the Mantel-Cox (log-rank) test. Data are presented as the mean ± standard error of the mean (SEM). Statistical significance was indicated as follows: ****P < 0.001, ***P < 0.005, **P < 0.01, and *P < 0.05. Unless mentioned otherwise in figure legends, experiments included a minimum of three biological replicates and were repeated independently at least three times.

## Acknowledgements

The authors gratefully acknowledge the support of Dr. Satya Prakash Yadav (Medanta Hospital, Gurugram, Haryana, India), Dr. Satyendra Katewa (SHALBY Sanar International Hospital, Gurugram, Haryana, India), and Dr. Manas Kalra (Sir Ganga Ram Hospital, Delhi, India) for their assistance in arranging the flow cytometry data of ALL patients. We also thank Dr Garima Nirmal for her assistance in obtaining the patient samples. We are deeply thankful to Professor Avinash Bajaj and his research team at the Regional Centre for Biotechnology, Faridabad, Haryana, India, for their contributions to the initial animal studies; the staff of CSIR–Indian Institute of Chemical Biology (CSIR-IICB) for their essential support in the pre-clinical studies; and the team at Adgyl Lifesciences, Hyderabad, India, for their help with animal studies. Special thanks are extended to Dr. Anurag Agrawal (Former Director, CSIR-IGIB) for his encouragement and support during the early phases of the project at CSIR-IGIB and the present Director of CSIR-IGIB, Dr. Souvik Maiti. The authors also appreciate the valuable contributions of Ankita Rajput, Aswathi C. B., Deeksha Joshi, Jahnvi Hora, Aparna Ramachandran, Syed Ahmad, and Dr. Suresh Madheswaran during the preparation of the manuscript. A patent application related to this work has been filed by Cellogen Therapeutics. We appreciate the support of Optical Microscopy Facility of the Advanced Technology Platform Centre (ATPC), Regional Centre for Biotechnology (RCB) for help with super resolution imaging.

---

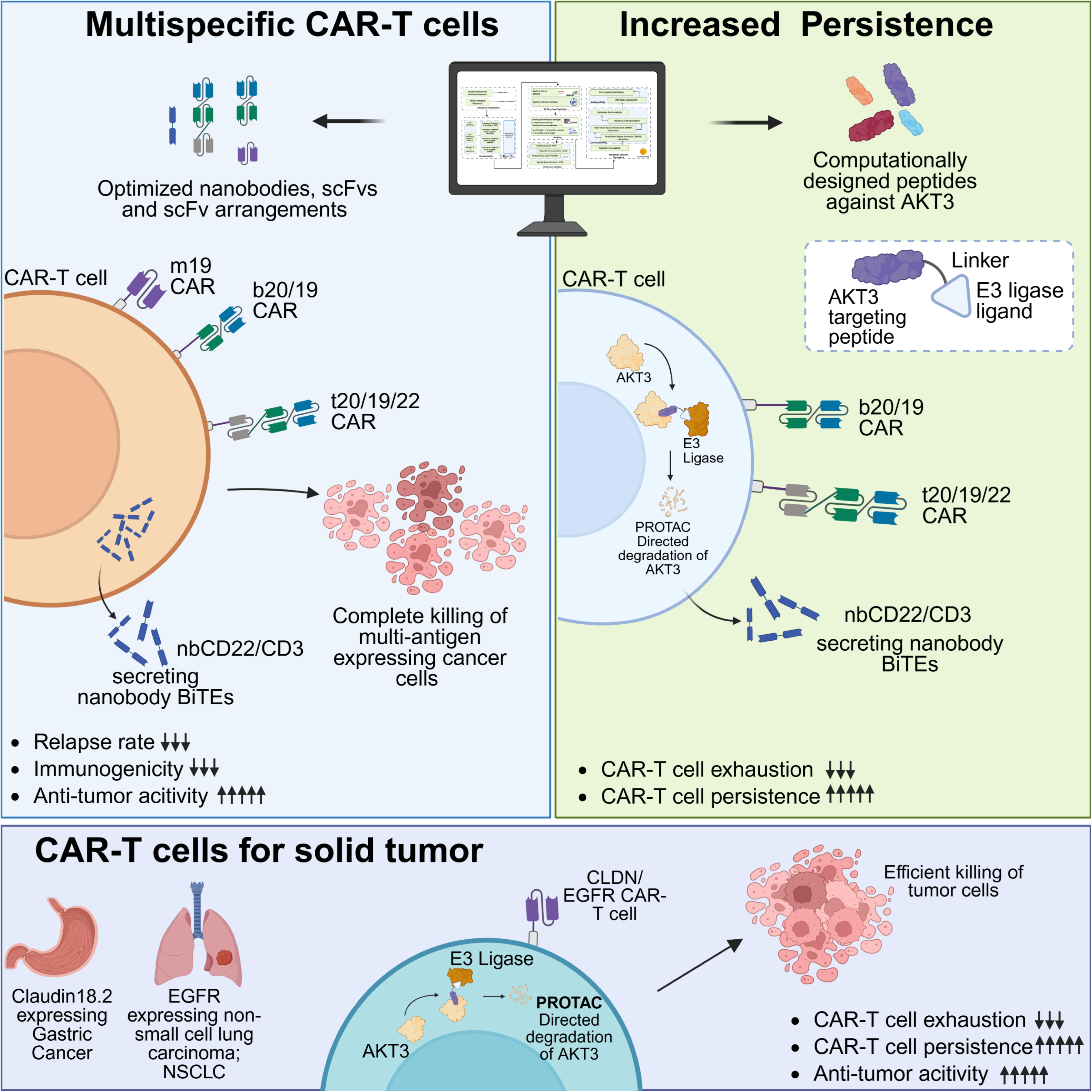

